# Forest structure, timber species regeneration, and timber volume dynamics along a logging gradient in a lowland tropical rainforest in Africa: Implications for biodiversity conservation and sustainable timber management

**DOI:** 10.1101/2025.03.26.645609

**Authors:** David Ocama Kissa, Emmanuel Fred Nzunda, Mnason Tweheyo, Daniel Lussetti, Enock Ssekuubwa, Douglas Sheil

## Abstract

Timber production, cutting and extraction is impacting vast areas of tropical forests, highlighting the need for management strategies to promote sustainable recovery of logged forests. However, limited information is available on how logging and enrichment planting affect forest structure, commercial tree species, and timber volume recovery. In this study, we assessed the effects of timber cutting and extraction (“logging”) on forest structure, regeneration of key timber species, and volume recovery across different logging intensities. We compared the effects of enrichment planting, initiated over 60 years ago, versus natural regeneration on timber volume recovery of *Khaya anthotheca* (Welw.) C.DC., a highly targeted species. We inventoried all live stems with a diameter at breast height (DBH) of ≥2 cm across 9 compartments using 45 plots of 0.5 ha each: heavily logged forests (25 plots, totaling 12.5 ha), lightly logged forests (15 plots, totaling 7.5 ha), and unlogged forest (5 plots, totaling 2.5 ha). Our results suggest that timber production has impacted on tree regeneration of harvested timber species such Entandrophragma species, *Milicia excelsa* (Welw.) C. Berg, *Olea capensis* L. subsp. welwitschii (Knobl.) Friis & Green and *Ricinodendron heudelotii* (Baill.) Heckel; Timber volume recovery of harvested species was 61.3% lower in heavily logged and 51.7% lower in lightly logged forests compared to unlogged forest. Stem density, basal area, and timber volume recovery of mahogany species were influenced by time since last logging. Notably, heavily logged forest that were enriched with *K. anthotheca* had significantly higher stem density and timber volumes of this species compared to logged forests without such enrichment. In conclusion, our study suggests that passive regeneration in Budongo’s logged compartments has been inadequate for achieving timber volume recovery of high-value commercial species. If an economic assessment proves favorable we recommend further trials of enrichment planting of high-value timber species (Mahogany spp., *Milicia excelsa*, and *Olea capensis*) alongside fast-growing species (*Maesopsis eminii* Engl., *Albizia* spp.). As well as Reduced Impact Logging and broader species selection from classes II and III, to reduce damage and enhance timber yields in production forests.

## Introduction

Tropical forests play an essential role in providing ecosystem goods and services [1]. They are important to the timber industry, contributing 24% of the USD 248 billion global forest product trade in 2019 [1]. In Uganda, sawn timber supply from tropical forests constitutes 30% at the domestic level [2] while at the international level, the estimate is 40% [3]. Global demand for primary processed wood products is projected to increase by 60% from 2020 to 2050 [4], while timber demand is expected to rise by 30% during the same period [5]. This increasing demand for processed wood products and for timber raises concerns about fulfilling future demand [6]. In Africa, the timber trade provides income to both local and national economies [6,7]. The global increase in demand for valuable hardwood timber has resulted in depletion of targeted timber species leading to degradation of large areas of forests [1,8]. Most timber from tropical forest countries is illegally logged, accounting for up to 30% of global timber production and 50–90% of harvesting in many tropical nations [9], including Uganda [10]. Illegal logging often causes greater forest damage than legal logging due to the lack of mitigation measures [11]. Yet, well-managed logged forests can retain significant conservation values [12] and offer safeguards for biodiversity thus contributing to nature protection [13,14].

Much as studies in tropical, subtropical and temperate forests have shown logging of trees for timber can enhance biodiversity and future timber production under appropriate post-harvest silvicultural practices and control of illegal activities [14–16]; there is still inadequate information on timber species and volume recovery [17]. Hence, the need to conduct studies to provide information to support sustainable management of logged forests [18]. Looking at recent studies that have examined the impact of logging on forest structure, timber species and volume recovery, most of them where concentrated on tropical forests outside Africa, e.g. in Asia [19,20]; Southern America [21–24]; Australia [25]; and few in Africa [26,27]. Although the studies revealed timber logging had negative effects on forest structure [20,28,29]; timber species recovery [30,31] and volume recovery [32,33], logging affects recruitment of timber species differently across ecological landscapes and logging intensities [34,35] and thus, there is need to evaluate timber species and volume recovery overtime to inform management decisions [25]. Again, understanding the effect of forest structural variables on recruitment of juveniles of timber species can generate information about the ecological mechanisms driving the recovery of the timber species [30,36].

Forest structure encompasses the vertical and horizontal distribution of aboveground vegetation resulting from forest dynamics, community demography and interactions [37,38]. Logging of large trees modifies forest structural attributes such as basal area, stem distribution, and stand density that may result into complex ecological niches for biological communities [15,39]. Hence, forest structure plays crucial role in understanding forest dynamics, timber yield and regulation of ecosystem services [40] as a result of its influence on tree recruitment [22] and species composition [36]. So, quantifying forest structural metrics e.g. tree (diameter, height, and basal area), canopy cover, undergrowth cover, and stem density [41] and their effects on tree juveniles can reveal the dynamics of the forest overtime [42] especially after logging of large trees. Logging of large trees changes canopy structure and understory vegetation, impacting tree recruitment, growth, and mortality across various size classes and growth forms [43]. Selective timber logging and high-intensity extraction can severely impact forest structure, hinder timber species regeneration, and reduce timber recovery [11]. Therefore, the combination of selective logging, extraction frequency, and recovery time since the last logging event can influence regeneration patterns, depending on the auto-ecology of the harvested species [44]. Justifying the need to conduct assessment of the impact of logging on understory/canopy structure, along with timber species regeneration and volume recovery so as to generate information for sustainable management of logged forests.

Many studies have assessed regeneration of timber species after logging with some indicating an increase, others a decrease, and some showing no effect at all [45,46]. However, the outcomes often depend on factors such as logging intensity, species-specific traits, and post-logging management. If not carefully planned, logging can lead to overexploitation of key commercial species, resulting in failed regeneration and poor timber volume recovery [47,48]. One study in the Amazon found that with a logging intensity of 20 m³/ha and a 40-year cycle, timber volume recovery may not reach pre-logging levels [49]. Despite an increase in merchantable species, continued extraction at 20 m³/ha every 30-40 years could lead to a decline in timber stocks in Amazonian production forests.

Timber production and ecosystem maintenance are key goals of sustainable forest management [50]. In tropical forests, reduced-impact logging minimises damage to residual trees and soil, promoting biodiversity and regeneration [29]. However, sustainable timber production remains challenging, particularly for high-value tropical hardwoods, due to regeneration failures even at low logging intensities, threatening long-term viability [49,51,52]. Thus, in areas with insufficient regeneration or depleted species, enrichment planting is vital, allowing the introduction of ecologically and economically important fast- and slow-growing species into degraded forests to restore composition and accelerate recovery [53,54]. For example, in the Amazon, enrichment planting with *Swietenia macrophylla* (American mahogany) and *Handroanthus serratifolius* significantly enhanced timber yields, with positive net value projections for a 60-year harvest cycle for *Swietenia* and a 90-year harvest cycle for *Handroanthus* [54]. Similarly, in Southeast Asia, trials with 22 locally adapted tree species found *Shorea macrophylla* and *Shorea ovalis* to have the highest internal rate of return (IRR) under low and high management costs, respectively, indicating the financial viability and ecological benefits of enrichment planting [55]. In Budongo forest, timber production was initiated through use of monocyclic and polycyclic logging methods [56]. Large sawmill operations were established between 1935 and 1992, with logging targeting trees of different size classes over time. From the 1940s to 1950, harvesting focused on trees with a DBH >130 cm, followed by trees with a DBH >80 cm from the 1960s to 1990, and finally, trees with a DBH >60 cm from 1991 to 2010, following a harvesting cycle of 60–80 years [56]. Timber volumes extracted from the compartments varied based on the availability of preferred resources, ranging from 30 to 60 m³/ha [57]. Post-harvest silvicultural operations such as enrichment planting of *Khaya anthotheca* (African mahogany)*, Entandrophragma angolense,* and *Milicia excelsa* (Welw.) C.C. Berg and poisoning of unwanted trees such as *Cynometra alexandri* C. H. Wright using arboricide chemical were implemented in 1950s-1960s so as to improve timber production in the logged compartments [56]. Although the survival of planted seedlings was lower than natural regeneration, partly due to damage by elephants, buffalos, and bush pigs among others, some compartments were well enriched with *Khaya anthotheca* [56]. To promote tree regeneration and sustain timber production, wildlife hunting in the 1960s led to the elimination of elephants and buffalos from Budongo Forest, driving them into Murchison National Park. Additionally, timber salvage through pit sawing was permitted in previously logged compartments from 1980 to 2010 [58]. However, due to declining timber resources, large sawmill operations ceased in 1992 [58], and pit sawing was banned in 2012 due to illegal activities and overexploitation [59]. Currently, there is limited information on forest dynamics, such as timber species and volume recovery after logging, which is essential for guiding sustainable management of logged forests [60]. Information on timber species and volume recovery after logging and or silvicultural operations [61] can help forest managers design effective restoration strategies to enhance timber production and maintain ecosystem services [62].

This study aimed to assess changes in forest structure, commercial timber regeneration, and timber volume recovery across different logging intensities. The specific objectives were to evaluate forest structure (vertical stratification, basal area, stem density, canopy cover, dominant heights, and undergrowth cover), the regeneration status of commercial timber species, and the recovery of merchantable timber volume. We hypothesised that:

a) Increasing logging intensity reduces stem density and basal area while altering vertical stratification, with potential long-term effects on forest recovery.
b) Logging intensity influences tree regeneration and timber volume recovery, with heavily logged areas experiencing greater impacts;
c) Over the 60-year harvesting cycle prescribed for Budongo, the merchantable volume of harvested timber species is expected to recover to levels comparable to unlogged forests, with enriched compartments supporting higher *Khaya anthotheca* volumes than non-enriched ones.

## Materials and methods

### Description of the study area

The study was conducted in Budongo Forest, a lowland semi-deciduous moist tropical rainforest in western Uganda (1°37′–1°55′ N, 31°22′–31°46′ E) [58]. Budongo Forest is one of Uganda’s 506 central forest reserves managed by the National Forestry Authority (NFA). Field research was conducted under NFA permit No. **372**, issued on June 22, 2022. Budongo Forest was gazetted in 1932 and covers approximately 825 km² [63]. It serves as a key water catchment for the Nile and Lake Albert [64]. The forest lies at an average altitude of 1,100 m (range: 700–1,270 m) with flat to slightly undulating terrain [57]. It experiences a bimodal rainfall pattern, with an annual mean of 1,300 mm (range: 1,200–1,800 mm year⁻¹) and a mean temperature of 21°C (range: 17–29°C) [65]. The soils are classified as Lixisols with a near-neutral pH and a high base saturation [66]. The forest has majorly four types of vegetation i.e. 1) Cynometra forest-dominated by an Iron wood species (*Cynometra alexandri*); 2) Mixed forest-dominated by Gambeya species and *Khaya anthotheca*; 3) Colonizing forest-dominated mostly by *Meosopsis eminii* and *Olea welwitschii* and 4) Swamp forest-composed of riparian mixed forest found along streams and dominated by *Pseudospondias microcarpa* and *Mitragyna stipulosa* [67]. The floristic composition is similar to those of the great Ituri Forest of the Democratic Republic of Congo [65].

The forest has been under the influence of different management regimes with sporadic pitsawing from 1920s before introducing mechanical logging around 1935 [68]. Importantly, most forest compartments have undergone selective logging and arboricide treatment with exception of designated nature reserve compartments [57]. Overall, 13 commercial timber species were harvested i.e. 4 mahogany species (*Khaya anthotheca, Entandrophragama utile, E. cylindricum* and *E. angolense*) and 9 other species (*Milicia excelsa, Lovoa trichilioides* Harms, *Cynometra alexandri, Erythropyleum suaveolens* (Guill. & Perr.) Brenan*, Mildbraediodendron excelsum* Harms*; Maesopsis eminii, Alstonia boonei* De Wild., *Ricinodendron heudelotii,* and *Morus mesozygia* Stapf) [68]. Enrichment planting using mahogany species mostly was conducted to increase the timber stocks [58]. Concerns have arisen over the declining fruiting of timber species in Budongo Forest, with only 10 of 62 monitored *Khaya anthotheca* trees fruiting once in 20 years since 1997 [69]. This trend poses a threat to the future of timber production as the local communities around Budongo especially from the southern part depends heavily on the forest resources [69]. Currently, forest protection is a big challenge in Budongo as illegal logging especially of poles and timber are more pronounced in almost all the compartments. Our study was conducted in Budongo Forest as shown (Fig 1).

**Fig 1.**
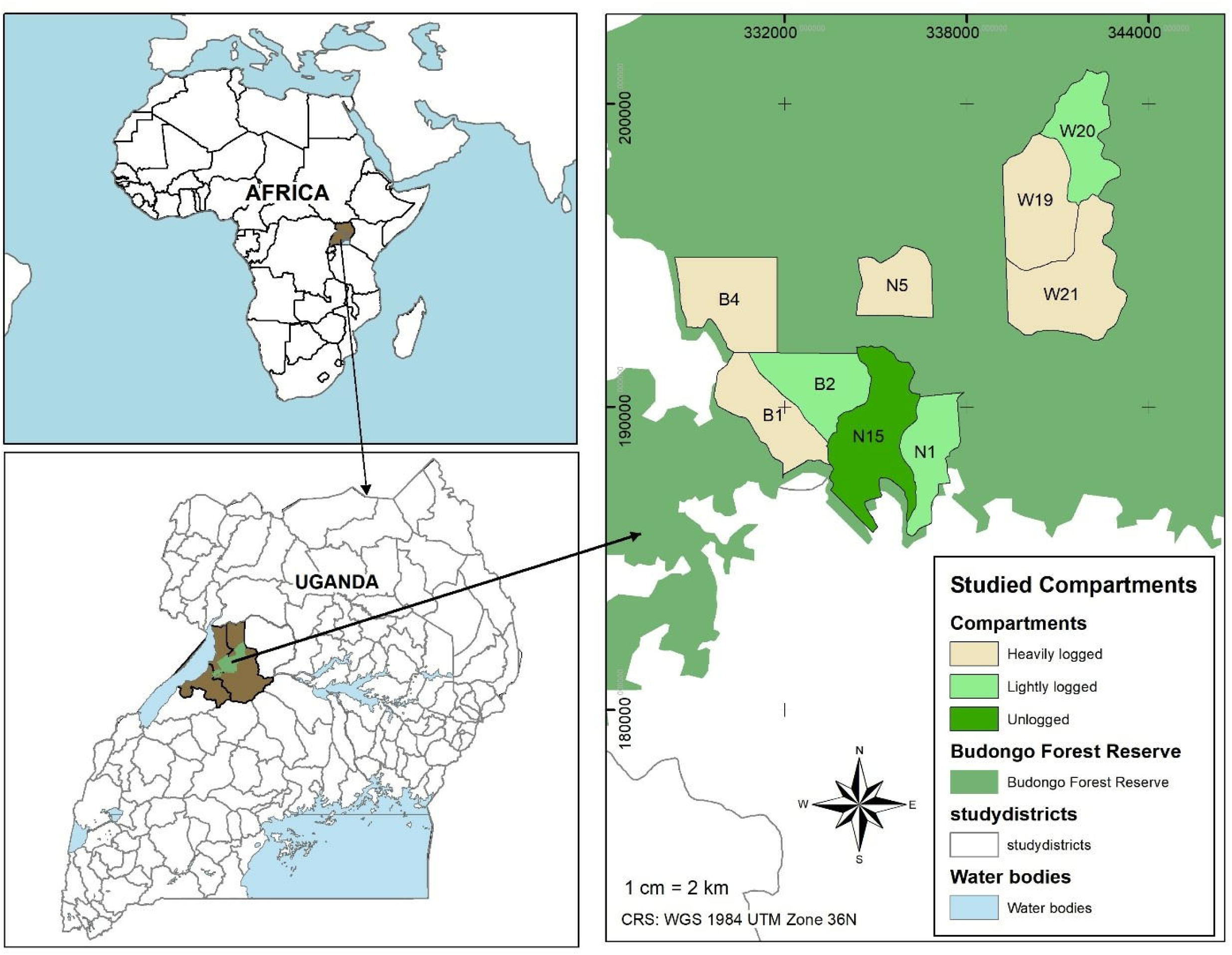
Map showing the study sites and logging gradient within Budongo Forest, East Africa. The Budongo Forest layer source is from [58].

### Study design

For this study, we selected nine compartments i.e. N1, N5 and N15 in Nyakafunjo block; B1, B2 and B4 in Biiso block; and W19, W20 and W21 in Waibira block which were logged at different time intervals to form the logging gradient (Table 1). We used every available information on logging records from the management plans to generate three logging intensities Kissa et al., in press). To analyze the effect of enrichment planting versus natural regeneration on timber volume recovery, we used four compartments: N1 (heavily logged, enriched with mahogany, and treated with arboricide), B2 and W20 (logged, treated with arboricide, and left to regenerate naturally), and N15 (a control, unlogged nature reserve with no arboricide treatment). We used a rectangular plot design of 100 m × 50 m, establishing five plots alternately at 200 m intervals along a 1-km transect to encompass different environmental and topographic gradients. Each 100 m × 50 m plot was subdivided into five 20 m × 50 m sub-plots to facilitate measurements and capture greater variability in gradients.

**Table 1.**
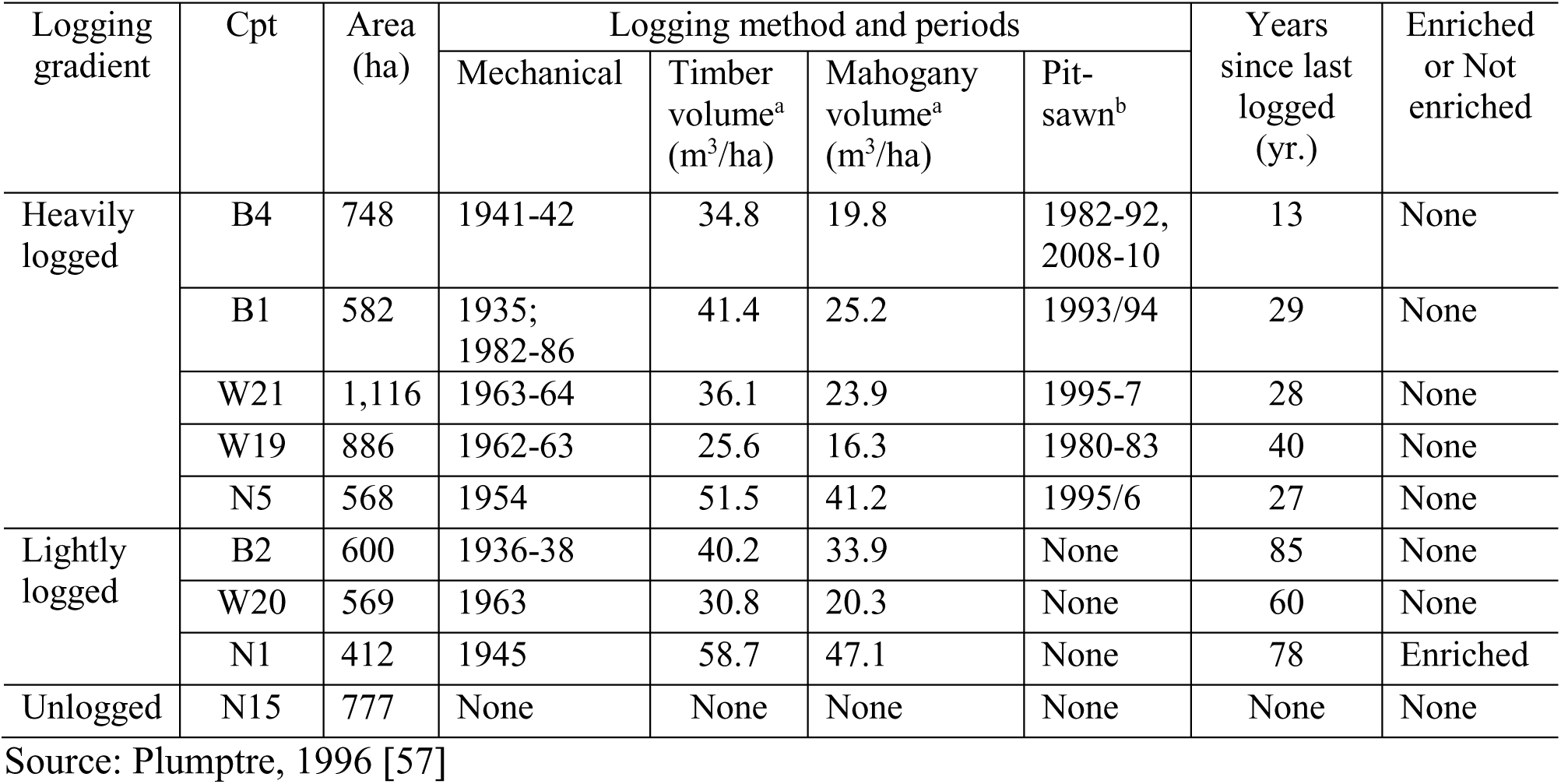
Logging intensities within the different compartments (Cpt) in Budongo forest, western Uganda. Timber volume^a^ = known harvested timber volume through mechanical method and Pit-sawn^b^ = unknown harvested timber volume through pitsawing

| Logging gradient | Cpt | Area (ha) | Logging method and periods |  |  |  | Years since last logged (yr.) | Enriched or Not enriched |
| --- | --- | --- | --- | --- | --- | --- | --- | --- |
|  |  |  | Mechanical | Timber volume<sup>a</sup> (m<sup>3</sup>/ha) | Mahogany volume<sup>a</sup> (m<sup>3</sup>/ha) | Pit-sawn<sup>b</sup> |  |  |
| Heavily logged | B4 | 748 | 1941-42 | 34.8 | 19.8 | 1982-92, 2008-10 | 13 | None |
|  | B1 | 582 | 1935; 1982-86 | 41.4 | 25.2 | 1993/94 | 29 | None |
|  | W21 | 1,116 | 1963-64 | 36.1 | 23.9 | 1995-7 | 28 | None |
|  | W19 | 886 | 1962-63 | 25.6 | 16.3 | 1980-83 | 40 | None |
|  | N5 | 568 | 1954 | 51.5 | 41.2 | 1995/6 | 27 | None |
| Lightly logged | B2 | 600 | 1936-38 | 40.2 | 33.9 | None | 85 | None |
|  | W20 | 569 | 1963 | 30.8 | 20.3 | None | 60 | None |
|  | N1 | 412 | 1945 | 58.7 | 47.1 | None | 78 | Enriched |
| Unlogged | N15 | 777 | None | None | None | None | None | None |

Source: Plumptre, 1996 [57]

### Field data collection

We conducted field tree inventory between November 2022 and December in heavily logged, lightly logged, and unlogged compartments (Figure 1). A total of 45 plots (100 m × 50 m) were sampled, covering 22.5 ha: 5 plots in unlogged, 15 in lightly logged, and 25 in heavily logged compartments. In each plot, we recorded all living trees, palms, and tree ferns with a diameter at breast height (DBH) ≥ 2 cm, along with cut stems (poles/timber) and snags [70] . Diameter was measured at 1.3 m above the ground (or above buttresses if necessary) using a diameter tape, and tree heights for stems with DBH ≥ 10 cm were measured with a Haglöf Vertex IV Hypsometer. Tree dominant heights were calculated from 50 mature, large-diameter live dominant trees. All trees were identified to species level by experienced parabotanists. Species names were confirmed based on literature [71,72]. Specimens of unidentified plants were collected and deposited at Makerere University herbarium.

At the center of each 20 m × 50 m sub-plot, we measured canopy closure in four cardinal directions (N, E, S, and W) using a Spherical Crown Densiometer [73]. The latitude and longitude of each plot center were recorded with a handheld Garmin GPSMAP 64s device. Undergrowth vegetation cover was assessed using a chequered board with 25 equally sized squares (10 × 10 cm), held at the plot center and observed from a distance of 10 m.

We categorized observations into four life forms based on height and diameter: treelets (height < 10 m; diameter < 10 cm), understory (height 10–20 m; diameter 10–30 cm), lower canopy (height 20–30 m; diameter 30–60 cm), and upper canopy (height > 30 m; diameter > 60 cm) [68]. This classification was informed by literature on the growth characteristics of indigenous Ugandan tree species [71,72]. Tree species were further grouped into juveniles (DBH < 10 cm) and established trees/shrubs (DBH ≥ 10 cm) [74]. Historically, tree species were classified as commercial or non-commercial based on key wood properties, including strength, durability, and the production of large, clear logs for lumber and veneer [75]. In modern contexts, high-value timber is predominantly defined by its economic market value, driven by wood durability and market demand [76]. Globally, a working list of commercial timber species has been established to guide international trade [77]. In Uganda, hardwood timber species are broadly categorized into three groups: high value, medium value, and low value traditional species [10]. In Budongo Forest, tree species are classified as commercial or non-commercial, with timber further graded into Classes I (high value), II (medium value), and III/IV (low value) based on strength, durability, and market preferences [58,78]. From Uganda’s Class I (high value), we selected key commercial timber species for their high market value and overexploitation. We anticipate the list will change to accommodate other species, as future classifications may incorporate lesser-known species that gain economic significance due to emerging market demand. To analyze the effects of logging intensities on timber species and volume recovery, stem diameters were classified into three groups: small/juveniles (DBH < 10 cm), medium trees (DBH 10–60 cm), and very large/mature trees (DBH ≥ 60 cm) [79].

### Statistical analysis

#### Forest structure (stem density and basal) along logging gradient

We compared stem density, basal area, canopy cover, and dominant heights across heavily logged, lightly logged, and unlogged forests. Stem density (N ha⁻¹) and basal area (m² ha⁻¹) were calculated for different tree categories using the formula proposed by [70].

We used descriptive statistics, one-way ANOVA and post-hoc Tukey’s Honest Significant Difference (HSD) test to compare for any significance difference (P-value was set at < 0.05) among the logging intensities [70]. Homogeneity of variance was tested using Levene’s test. For data sets that didn’t conform to ANOVA test of normality, we used Welch’s ANOVA followed by Games-Howell post hoc test due to unequal sample sizes. We further analysed the effect of recovery time on stem density and basal area recovery of small, medium and large tree category using a GLM as indicated in equation 1 and 2 [80]. All analysis were done were using R Software [81].

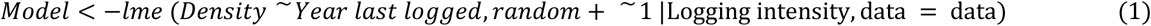

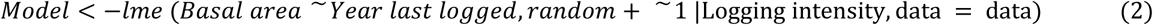

#### Regeneration status

Regeneration status was classified as *good, fair, poor,* and *none* based on the abundance of juveniles (saplings and poles, DBH ≥2 cm to <10 cm) relative to adults (DBH ≥10 cm) across all sites within a logging intensity. A tree population is considered to be of *good* regeneration status if their juveniles outnumbered adults, *fair* regeneration status when juvenile and adult populations were equal, *poor* regeneration when adults were predominant with few juveniles, and *none* when a tree species was present only in the adult stage without juveniles [82,83].

#### Relationships between forest structural parameters (basal area, stem density and canopy cover) and abundance of juveniles (all timber species, mahogany species and other harvested timber species) along logging intensity

The relationship between forest structural parameters and juvenile (saplings and poles) of all tree species, timber species and mahogany species) were analysed using Generalized Linear Mixed-Effects Model (GLMM) with a Poison or binomial distribution and logit link. Juvenile density was fitted with GLMM models using a Poisson distribution and log link. We used *“lme4”* package for maximum-likelihood estimation of the parameters of the GLMM models [84]. The goodness of fit of the models were compared using Akaike’s information criterion (AIC) [85]. We included plots as random effect, forest structural attributes and abundance of juveniles as fixed effects and logging intensity as interactions. The equation is shown below;

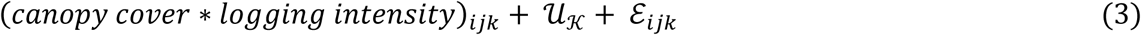

where Juvenile abundance _ijk_ is the abundance of juveniles in plot I, within logging intensity j (e.g. heavily logged, lightly logged and unlogged); ijk is the expected juvenile abundance with a log link function applied; ß_0_ is the intercept representing the baseline abundance when all predicators are zero; ß_1’_ ß_2;_ ß_3_ and ß_4_ are coefficients for basal area, stem density, canopy cover and logging intensity. Logging intensity is treated as categorical variable; ß_5,_ ß_6_ and ß_7_ are the coefficients for the interactions between basal area and logging intensity, stem density and logging intensity and canopy cover and logging intensity; U_k_ is the random intercept for plot to account for variability among plots; εij is the residual error term.

We used a GLMM to analyse forest structural parameters (stem density, basal area, and volume) across logging intensities, with time as a measure of successional recovery since logging simplifies the age structure of the forests (Edwards et al., 2019). We included logging intensity (categorical) and years since the last-logged (continuous) as fixed effects, while plots were random effect. This approach provided insights into vegetation changes and forest dynamics across different logging intensities and recovery stages. We tested the normality with the Shapiro-Wilk test and the analyses of GLMs were performed with the *“nlme”* package [86]. We grouped trees into timber and non-timber species, juveniles, medium and large trees as categorical variables and tested if years since last logged has an effect on variation in density and basal area. All analysis were done were using R Software [81].

#### Estimation of timber volumes for the different logging gradient

We estimated the merchantable Volume (V_m_) using the formula developed by [68] for Budongo as :

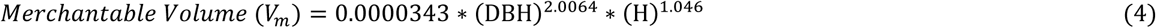

Where V_m_ is the estimated merchantable volume (m^3^); DBH is the diameter (cm) and H (m) is the height from the ground to the smallest diameter immediately below the insertion of the first major live branch.

Generalized linear mixed-effects model were used to determine the effect of time since last logging took place on timber volume recovery (all timber species, mahogany species and other harvested timber species) [85,86]. Timber volume and year since last logged were treated as fixed effects and logging intensity included as random effect into the model shown below;

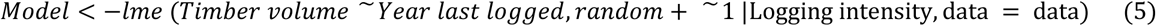

We compared the timber volume of *Khaya anthotheca* between compartment N1 which was logged and enriched in 1960s with those compartments (W20 and B2) which were logged over 60 years but not enriched. We included compartment N15 as a control since no official logging has occurred to date, and it has remained a strict nature reserve. Given that mechanised logging began in Budongo in 1935, we assumed N15 has remained unlogged for 88 years. We standardised annual recovery by normalizing it with the volume removed from each compartment, ensuring comparability, as outlined in Equation 6. We used generalized linear mixed-effects model to assess the effect of enrichment on the *K. anthotheca* timber volume using the equation 7.

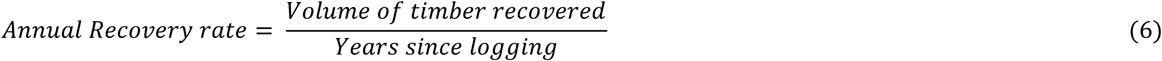

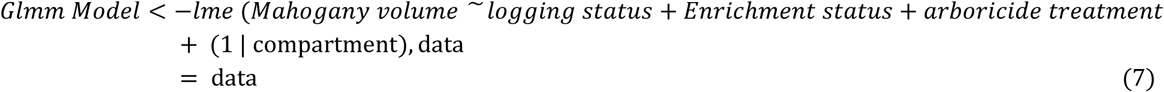

Mahogany volume was estimated for each plot surveyed within each compartment. Each compartment was categorized as (a) unlogged or (b) logged, with enrichment recorded as yes or no, and the amount of arboricide applied, as reported by [57]. However, the specific arboricide quantity for compartment N1 was not provided. To address this, we applied arboricide treatment patterns from the two neighboring compartments and conducted a sensitivity analysis to assess the impact of varying arboricide levels on mahogany volume recovery. All the data were tested for normality with the Shapiro-Wilk test before the analyses were performed with the package *“nlme”* [86]. All analysis were done were using R Software [70].

## RESULTS

Multi-stem trees (1.3%) were counted as individual trees, yielding a total of 37,582 live individual trees comprising 457 individuals (1.2%) of treelet species, 20,029 individuals (53.3%) of understory species, 5,068 (13.5%) lower canopy species, and 12,028 (32%) upper canopy species.

Among all the individuals, 17,031 (45.3%) were commercial timber species while 20,551 (54.7%) were non-commercial (S1). Out of the 17,031 commercial stems, 1,382 (8.1%) were tree species categorized as class I due to their high valuable timber (1,122 individuals of mahogany species-*Khaya anthotheca*; *K. grandifoliola*; *Entandrophragama utile*; *E. angolense*; *E. cylindricum* and 260 individuals of other class I timber species-*Milicia exc*elsa; *Lovoa trichilioides*; *Olea capensis*; *Leplaea cedrata*;*Zanthoxylum gilletii*) while the remaining individuals i.e. 15,649 (91.9%) were timber tree species of class II and III/IV categories.

### Variation of forest structure (stem density and basal area) for different growth-forms along the logging gradient

Our results showed that stem density and basal area (DBH ≥2 cm) varied across growth forms and size classes (Appendix Table 2). One-way ANOVA revealed significant variation in stem density among logging intensities (F₂, ₄₂ = 11.982, p < 0.001) and interaction between logging intensity and growth form (F_6, 156_ = 6.062, p < 0.001). Logged forests had lower understory densities (793.6 ± 459.7 stems/ha in heavily logged, 826.1 ± 247.6 stems/ha in lightly logged, and 1,565.2 ± 258.5 stems/ha in unlogged) and canopy tree densities (648.24 ± 196.46 in heavily logged, 871.20 ± 182.47 in lightly logged, and 1,070.80 ± 201.32 in unlogged) (S3). One-way ANOVA revealed significant variation in stem density among logging intensities for understory (F₂, ₄₂ = 8.719, p = 0.001) and canopy trees (F₂, ₄₂ = 3.251, p = 0.043). Tukey HSD post-hoc tests showed understory tree density differed significantly between unlogged and heavily logged (p < 0.001) and unlogged and lightly logged forests (p ≤ 0.001), but not between lightly and heavily logged (p = 0.490).

Canopy tree density differed only between unlogged and heavily logged forests (p = 0.013), with no significant difference between unlogged and lightly logged (p = 0.078) or lightly and heavily logged (p = 0.335). Understory vegetation (DBH <10 cm) was dominated by four species i.e. *Drypetes ugandensis*, *Lasiodiscus pervillei*, *Celtis mildbraedii*, and *Rinorea beniensis* which accounted for 45.5% of total stem density and were most abundant in unlogged forests. For instance, *D. ugandensis* had 150.4 stems/ha in unlogged forests but only 6.4 and 2.9 stems/ha in lightly and heavily logged forests, respectively. Similarly, *L. pervillei* had 654.8 stems/ha in unlogged forests, compared to 324.0 and 248.6 stems/ha in lightly and heavily logged forests. *C. mildbraedii* had 382.0 stems/ha in unlogged forests, decreasing to 149.9 and 174.6 stems/ha in lightly and heavily logged forests, while *R. beniensis* had 294.4 stems/ha in unlogged forests, compared to 173.2 and 252.7 stems/ha in logged forests. The high stem density of these late-succession species suggests greater stem packing in unlogged forests, where they persist in the understory, awaiting favorable conditions for growth. We observed a high occurrence of *R. beniensis* in plots with abundant large *Cynometra alexandrii* trees, indicating a possible species association.

The basal area data reveal significant impacts of logging intensity on various growth forms (Table 2). Unlogged compartments consistently show higher basal area in treelets, understory, and upper canopy species compared to both lightly and heavily logged areas. Heavily logged compartments, in particular, exhibit the lowest basal area across treelets and understory species, indicating severe disruption to regeneration. Lightly logged areas show intermediate basal areas, suggesting they allow some recovery. The lower canopy appears more resilient, with no significant differences observed in basal area across logging intensities. Mature trees in the understory and upper canopy also show lower basal areas in logged areas, with unlogged compartments maintaining higher values. Overall, logging, especially heavy logging, significantly reduces basal area and disrupts regeneration, particularly for treelets and understory species, but some growth forms like the lower canopy may be more resilient.

We assessed the influence of time since last logging on the density recovery of small (DBH <10 cm), medium (DBH ≥10 to <60 cm), and large stems (DBH ≥60 cm), as well as understory and canopy trees (S3). GLMM results indicated a significant effect of time since logging on the density of small, large, and canopy trees, while no significant effect was observed for medium and understory trees (Fig. 2A-E).

**Fig. 2.**
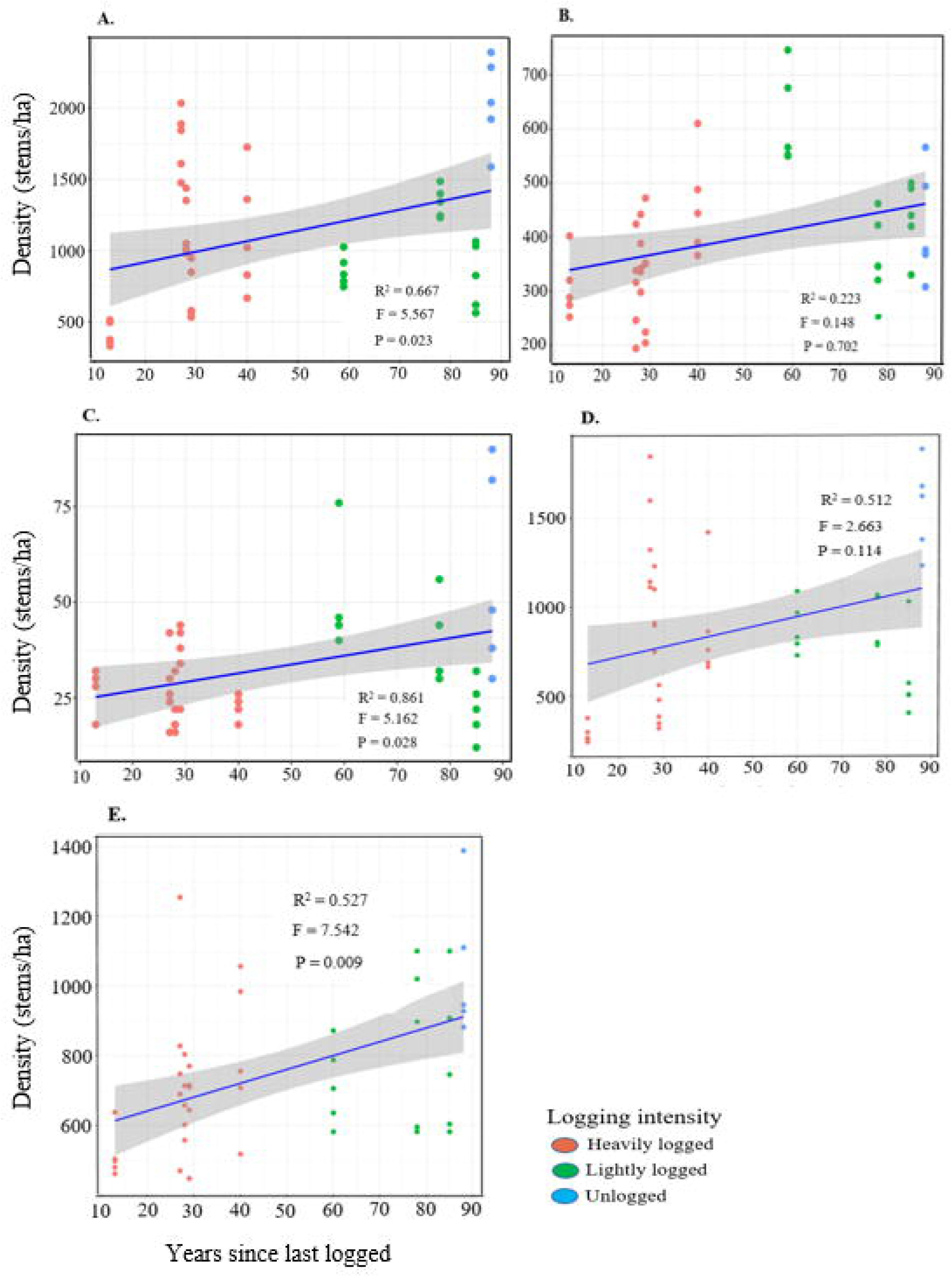
Effect of time since last logging on the density of all trees across different diameter categories: A) small, B) medium, C) large trees, D) understory trees and E) canopy trees in Budongo Forest. Density of trees in each plot is represented by a single dot. The R^2^ is included to explain the fixed and random effects of the model. The F and P-values are the summary of the significance of the model.

### Variation of forest structure (stem density and basal area) for all timber species, mahogany species and other harvested species along the logging gradient

Stem densities vary significantly across logging intensities for all tree categories, including timber species, mahogany, and other harvested species (Table 3; raw dataset in S4). One-way ANOVA showed significant variation in stem density for small (DBH <10 cm, p = 0.002) and large trees (DBH ≥60 cm, p < 0.001), but not for medium trees (DBH ≥10 to <60 cm, p = 0.514). Tukey HSD post-hoc tests confirmed differences among logging intensities for small and large trees.

**Table 3.**
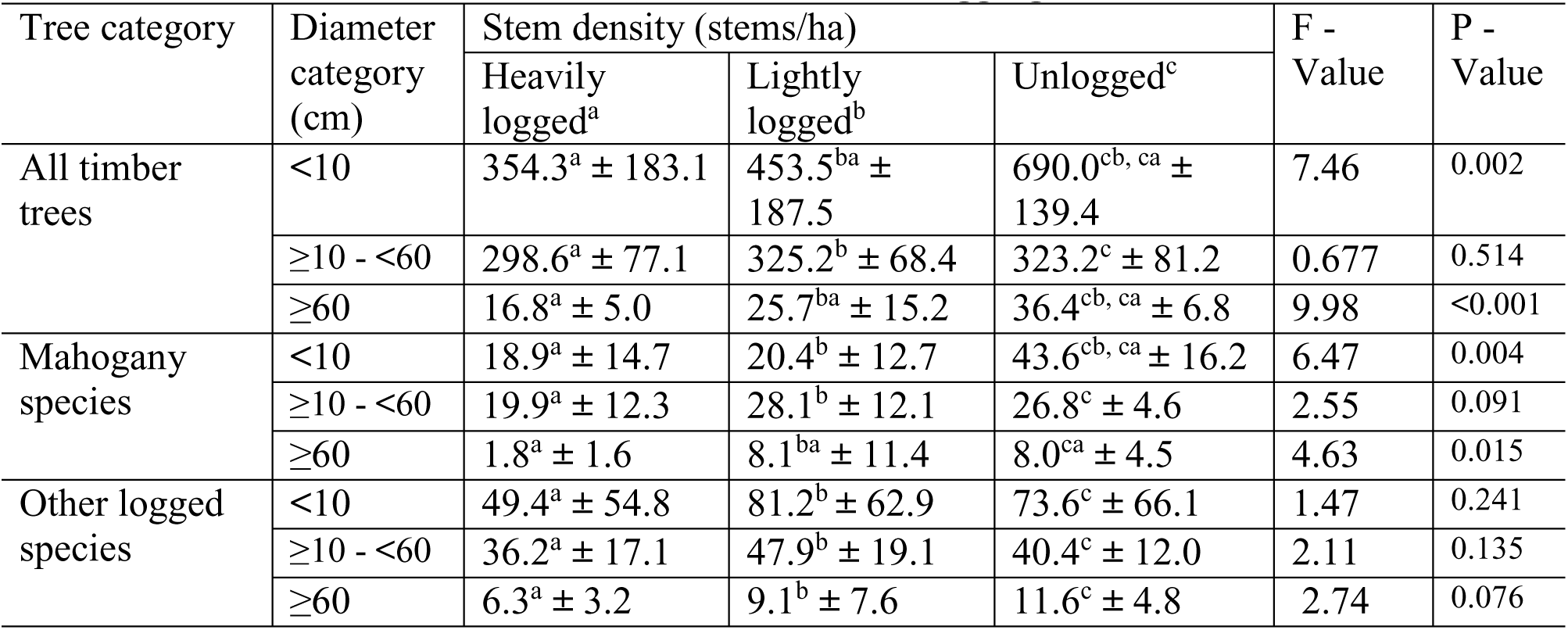
Summary of forest structure, presenting stem density (stems/ha ± SD) for all timber species, mahogany species, and other harvested species by diameter categories (DBH <10 cm, DBH ≥10 to <60 cm, and DBH ≥60 cm) under different logging intensities.

Unlogged forests had significantly higher stem densities across most diameter classes compared to lightly and heavily logged forests. The difference was most pronounced for small trees, where unlogged forests had nearly twice the density of heavily logged forests (690.0 ± 139.4 vs. 354.3 ± 183.1 stems/ha, p = 0.002). Similarly, large-diameter trees (≥60 cm DBH) were more abundant in unlogged forests (36.4 ± 6.8 stems/ha) than in lightly logged (25.7 ± 15.2 stems/ha) and heavily logged forests (16.8 ± 5.0 stems/ha, p < 0.001), indicating the long-term impact of logging on forest structure.

Among mahogany species, significant variation in stem density was observed for small and large trees (Table 3). Unlogged forests had higher densities of small mahogany trees than lightly and heavily logged forests. Likewise, large mahogany trees were more abundant in unlogged and lightly logged forests (8.1 ± 11.4 stems/ha) than in heavily logged forests (1.8 ± 1.6 stems/ha, p = 0.015). For other harvested timber species, no significant differences in stem densities were observed across logging intensities for any diameter class (Table 3), suggesting greater resilience to logging disturbances compared to mahogany and other high-value timber species.

Overall, these findings highlight the lasting effects of logging on forest composition and structure, particularly for commercially valuable species like mahogany. The significant reduction in large-diameter trees in heavily logged forests suggests long-term impacts on stand structure and species recovery.

Results of basal area for the different tree categories are indicated in (Table 3). Comparison of basal area using one-way ANOVA showed significant difference in tree basal area across logging intensities (F = 370.55, P < 0.0001). Tukey HSD post-hoc test showed significant difference in tree basal area between unlogged and heavily logged (P <0.001), unlogged and lightly logged (P < 0.001), and lightly logged and heavily logged (P <0.0001) for all the stem diameter categories. All timber species had significantly higher basal area in unlogged forests across all diameter categories, with the most pronounced differences in medium-sized (DBH ≥10 to <60 cm) and large trees (DBH ≥60 cm), where unlogged forests had the highest basal area, followed by lightly logged and heavily logged forests. Small trees (DBH <10 cm) also exhibited significant differences, with the highest basal area in unlogged forests (Table 4).

**Table 4.**
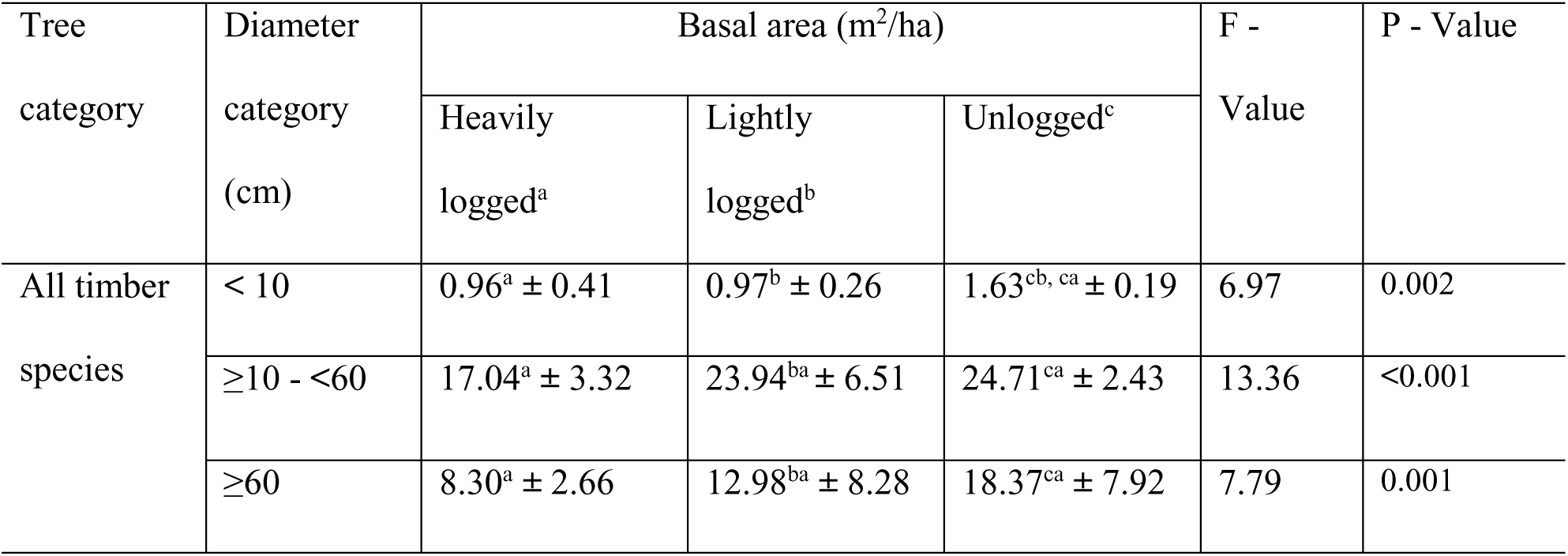
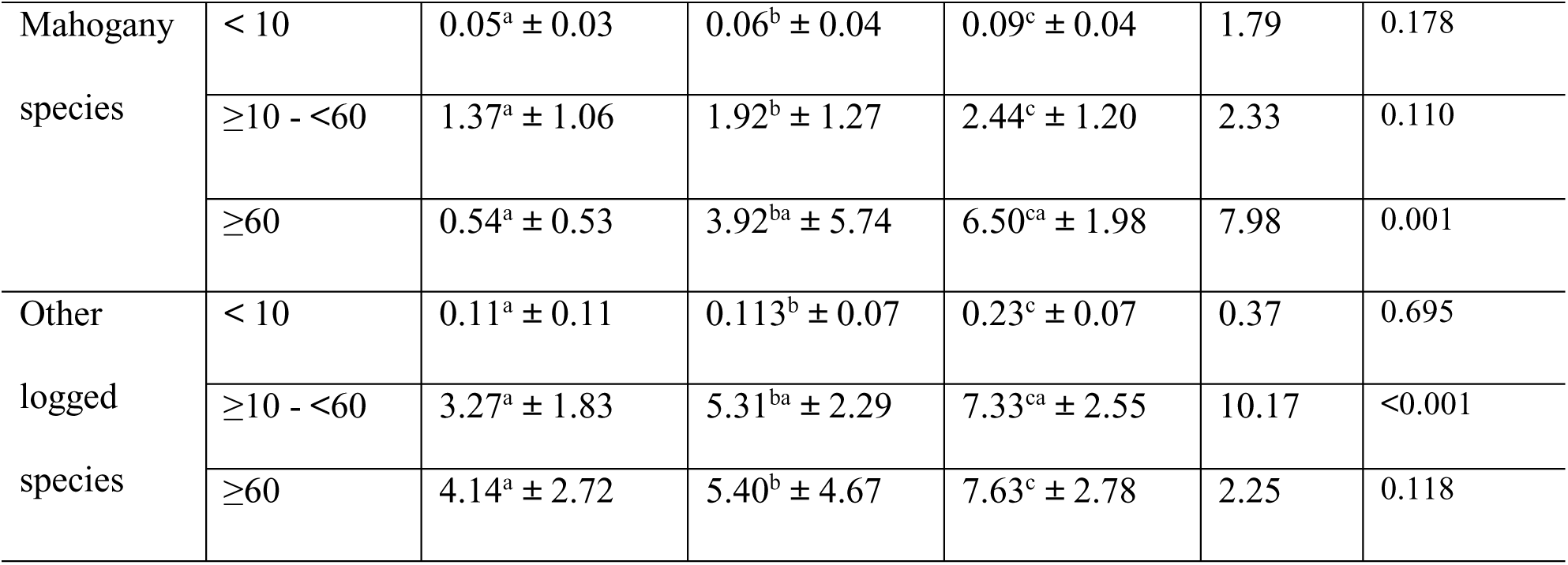
Summary of basal area (m²/ha ± SD) for all timber, mahogany, and other harvested species by diameter categories (DBH <10 cm, DBH ≥10 to <60 cm, and DBH ≥60 cm) under different logging intensities.

Mahogany species showed significant variation only in the large diameter class (DBH ≥60 cm), with basal area highest in unlogged forests (6.50 ± 1.98 m²/ha) and lowest in heavily logged forests (0.54 ± 0.53 m²/ha), indicating a substantial decline due to logging. No significant differences were observed for small and medium-sized mahogany trees.

Other harvested species exhibited significant variation in basal area for medium-sized trees (DBH ≥10 to <60 cm) (F₂, ₄₂ = 10.17, p < 0.001), with unlogged forests having the highest basal area. However, differences were not significant for small and large diameter classes (Table 4). These findings underscore the long-term impact of logging, with unlogged forests retaining higher basal area, especially for large trees. The significant decline in large mahogany trees in logged forests suggests high-value species are disproportionately affected.

### Variation in density of pioneers and late succession tree species along the logging gradient

The mean tree density (≥2 cm) of pioneer trees was 227.3 ± 157.9 stems/ha in heavily logged, 269.3 ± 133.2 stems/ha in lightly logged, and 217.5± 67.1 stems/ha in unlogged forests (S5). ANOVA indicated no significant difference in pioneer tree density among logging intensities (F₂, _42_ = 1.178, P = 0.318). In contrast, late-succession tree density was 1,294.2 ± 557.4 stems/ha in heavily logged, 1,347.1 ± 229.7 stems/ha in lightly logged, and 2,397.6 ± 273.4 stems/ha in unlogged forests. ANOVA revealed a significant difference in late-succession tree density across logging intensities (F₂, ₄₂ = 12.965, P < 0.001). Tukey HSD post-hoc tests showed significant differences between unlogged and heavily logged (P < 0.001) and between unlogged and lightly logged (P< 0.001), but not between lightly logged and heavily logged (P = 0.721).

### Variation in forest structure i.e. mean stem diameter, stand dominant heights, canopy closure, and undergrowth cover along the logging gradient

Results show that *Cynometra alexandri* had the largest stem diameter across all logging intensities, with a maximum DBH of 150.3 cm in unlogged forests, 148.5 cm in heavily logged forests, and 130.6 cm in lightly logged forests. The tallest tree was recorded in unlogged forests (42.4 m), followed by lightly logged (39.9 m) and heavily logged forests (37.3 m). Table 4 presents the forest structure results, including mean stem diameter, tree height, canopy cover, and undergrowth cover, based on dataset (S6). Comparison of mean differences using one-way ANOVA showed significant differences for all the variables between logging intensities (Table 5). Tukey HSD post-hoc test showed significant difference for all the variables across the logging intensity (Table5).

**Table 5.** Summary of forest structure i.e. mean stem diameter (cm ± SD) for trees with diameter (DBH ≥10 cm), mean tree height (m ± SD), mean canopy cover (%), and mean undergrowth cover (%) in each of the logging intensities.

| Category of forest structure | Logging intensity |  |  | F - Value | P - Value |
| --- | --- | --- | --- | --- | --- |
|  | Heavily logged | Lightly logged | Unlogged |  |  |
| Mean DBH ≥10 cm | 23.2<sup>a</sup> ± 1.9<br>(19.9 – 26.7) | 24.1<sup>b</sup> ± 1.7<br>(21.0 – 27.3) | 26.7<sup>cb, ca</sup> ± 1.9<br>(24.7 – 29.4) | 7.68 | 0.001 |
| Mean tree height (m) | 10.9<sup>ab, ac</sup> ± 0.8<br>(9.6 – 12.2) | 11.6<sup>ba, bc</sup> ± 0.7<br>(10.6 – 12.8) | 13.3<sup>ca, cb</sup> ± 0.6<br>(12.7 – 13.9) | 29.48 | <0.001 |
| Mean canopy height (m) | 17.6<sup>ab, ac</sup> ± 1.5<br>(15.2 – 20.9) | 19.8<sup>ba, bc</sup> ± 2.2<br>(16.9 – 24.0) | 24.2<sup>ca, cb</sup> ± 3.4<br>(20.5 – 28.0) | 24.61 | <0.001 |
| Canopy cover (%) | 75.2<sup>ba, ca</sup> ± 4.3<br>(67.9 – 82.0) | 84.1<sup>ba, cb</sup> ± 4.0<br>(74.7 – 89.0) | 89.5<sup>ca, cb</sup> ± 3.1<br>(84.8 – 93.3) | 37.67 | <0.001 |
| Undergrowth cover (%) | 71.0<sup>ab, ac</sup> ± 4.9<br>(62.7 – 79.1) | 59.8<sup>abb, ac</sup> ± 6.5<br>(48.0 – 68.9) | 47.6<sup>ac, bc</sup> ± 4.6<br>(41.8 – 54.2) | 47.01 | <0.001 |

± SD is the standard deviation, (minimum – Maximum values based on plots) for the different forest structural parameters.

### Regeneration status of harvested commercial timber species in Budongo Forest

Among the class I timber category (S7), only *K. anthotheca* had overall fair regeneration status while the rest showed poor regeneration with even less than one stem of juvenile per hectare in heavily logged compared to lightly logged or unlogged where the tree populations exists (Table 6). For class II timber category, only *C. alexandri* had overall good regeneration status while others have poor regeneration status in all the logging intensities. Tree species with none regeneration status were *O. carpensis* in class and *R. heudelotii* in class II within heavily logged forests. Overall, juvenile populations were very low for most tree species in heavily logged, followed by lightly logged and highest in unlogged. However, tree populations of these timber species i.e. *O. carpensis, L. trichilioides* and *M. excelsa* were not observed in unlogged forest.

**Table 6.**
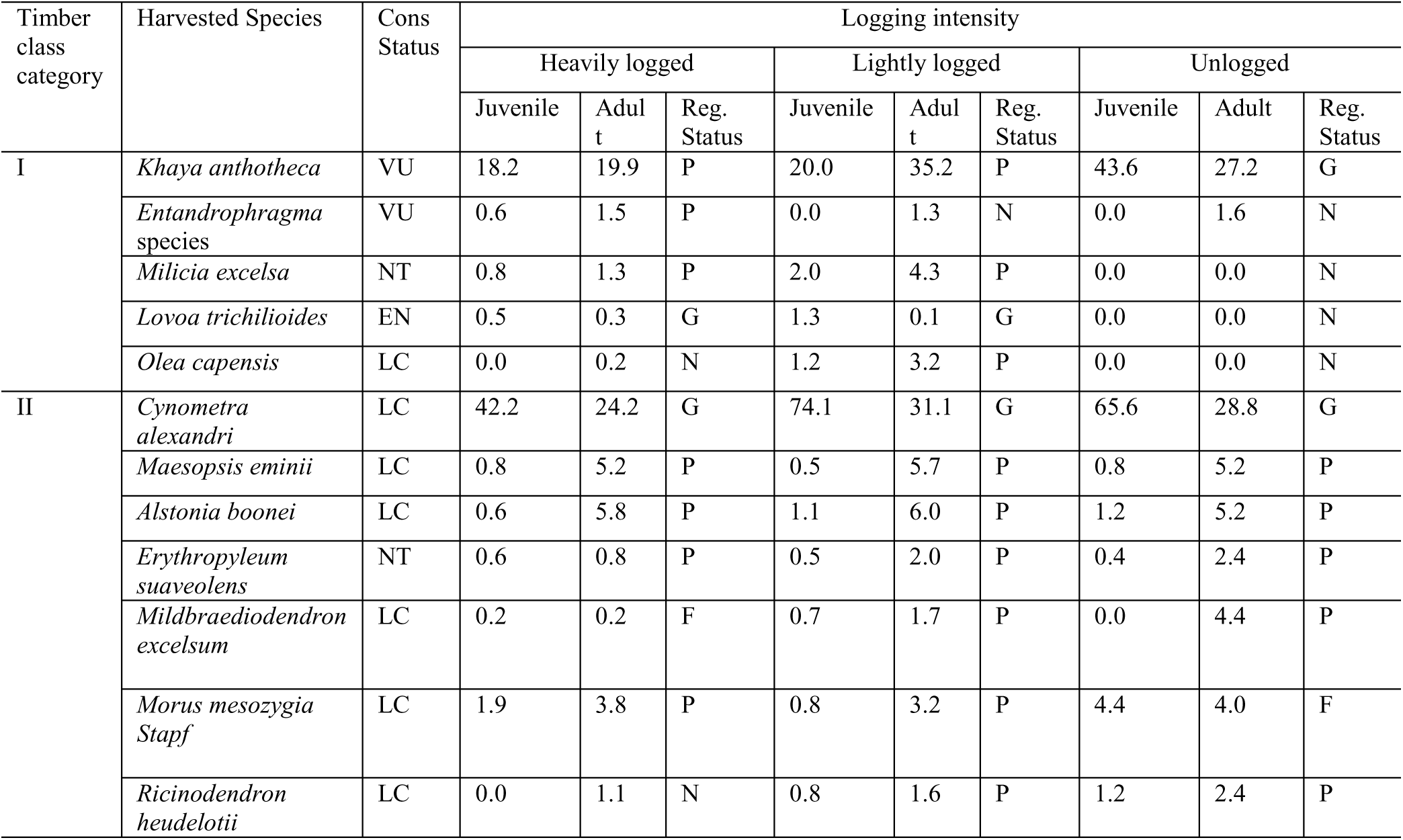
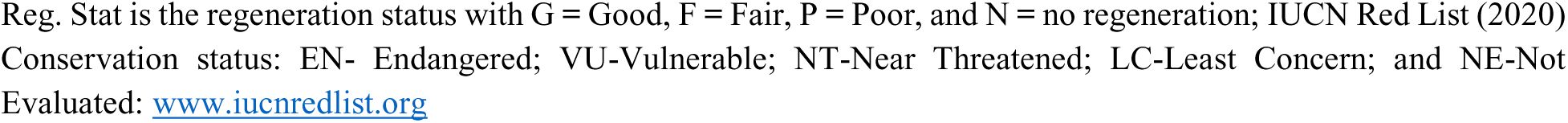
Regeneration status of harvested timber species based on the density of juveniles to adult populations (J: A) and their conservation in each logging intensity in Budongo forest.

### Abundance of class I, II, III/IV timber and non-commercial timber species for different diameter categories in each of the logging intensity

The proportion of tree species by timber class varies across logging intensities and diameter categories (Fig. 3), with the supporting dataset provided in Supplementary Material (S8). Class II species (red) dominated across all logging intensities and size classes, highlighting their resilience and suggesting intense selective harvesting of higher-value species. Class I species (blue) are more prevalent in unlogged forests, particularly in the largest diameter class (DBH ≥60 cm), but decline in logged forests, suggesting their preferential harvesting. Non-commercial species (yellow) are more abundant in smaller diameter classes (DBH <30 cm), especially in logged forests, implying a shift in species composition favoring non-commercial species post-logging. Class III/IV species (black) are relatively more common in the medium-sized category (DBH 30 to <60 cm), particularly in logged forests, indicating persistence at intermediate stages. Overall, unlogged forests retain a higher proportion of high-value Class I species, while logged forests show a decline in these species and an increase in non-commercial and lower-value timber species. This pattern highlights the long-term impact of logging, with regenerating forests being dominated by non-commercial species, potentially affecting future timber yields.

**Fig 3.**
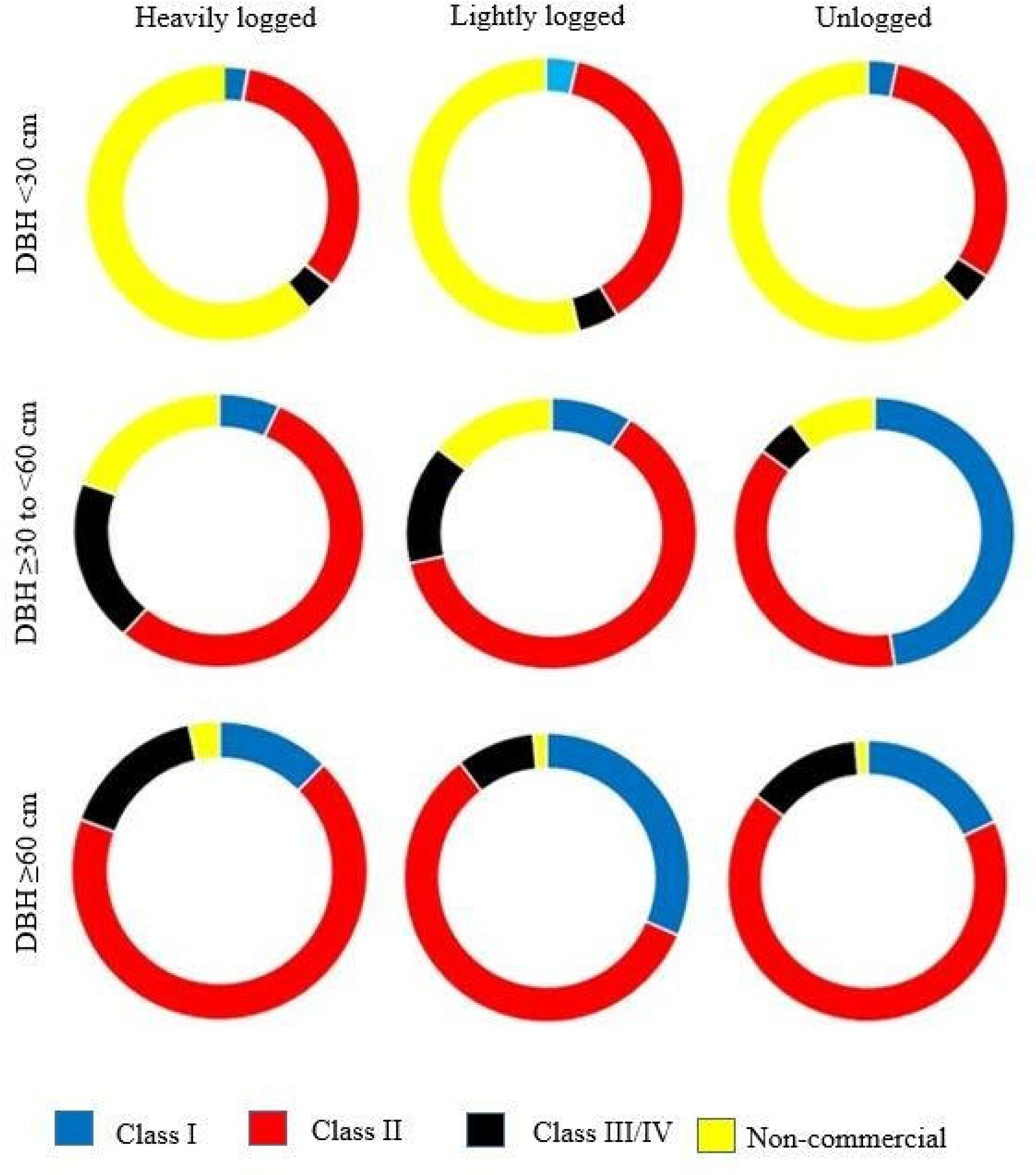
Proportion of tree species (commercial timber and non-commercial trees) by number of individuals for different diameter categories in each logging intensity in Budongo Forest.

### Relationships between forest structural parameters (stem density, basal area and canopy) and regeneration status (juvenile abundance) of (all timber species, Mahogany species and other harvested timber species) along logging intensity

We examined the effects of forest structure (basal area, stem density for DBH ≥30 cm, and canopy cover) and logging intensity on juvenile abundance of timber species, mahogany, and other harvested species (S9). GLMM results showed significant effects only for mahogany juveniles. Basal area (β = -2.335, P = 0.048), unlogged forest conditions (β = 412.436, P = 0.039), and interaction effects of basal area × logging intensity in lightly logged (β = 3.763, P = 0.025) and unlogged forests (β = 2.094, P = 0.023) were significant. Additionally, stem density × logging intensity interaction was significant in unlogged forests (β = -3.942, P = 0.036). Random effects analysis revealed substantial variation in juvenile density among plots (SD = 45.2), suggesting unaccounted environmental or ecological influences. These findings highlight basal area, logging intensity especially in unlogged and lightly logged forests, and plot-level variability as key drivers of mahogany juvenile density, while other factors had minimal impact.

We analysed the effect of time since last logging on juvenile abundance of all timber species, mahogany, and other harvested species (Fig 4). GLMM results revealed significant differences across all categories, including total timber species, mahogany, and other harvested species (Fig. 4A-C).

**Fig 4.**
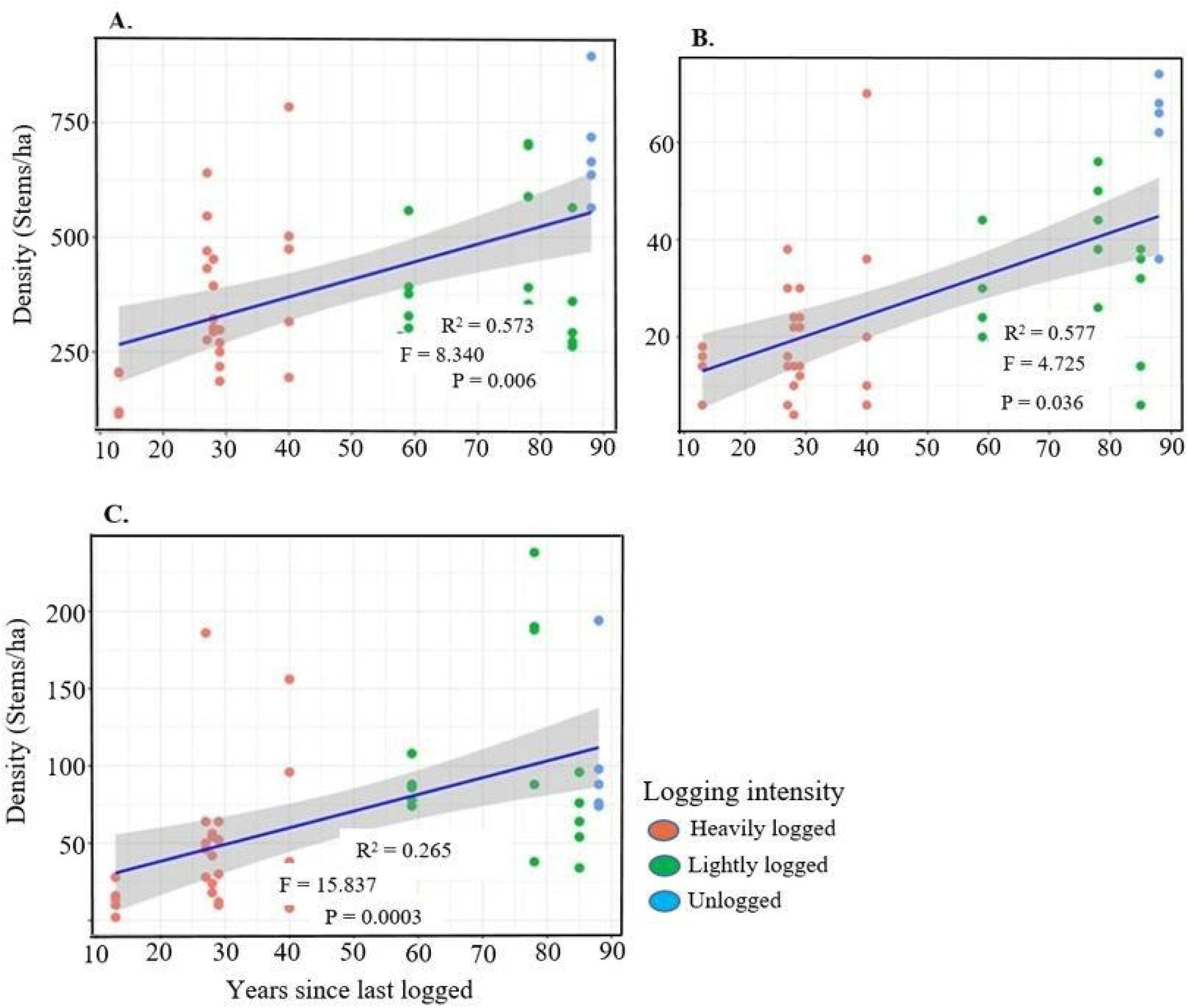
Effect of time since last logging on the juvenile abundance; A) all timber species, B) mahogany species, and C) other harvested timber species in Budongo Forest. Juvenile abundance in each plot is represented by a single dot. Plots with **red** color represent heavily logged areas, **green** indicates lightly logged compartments, and **blue** corresponds to plots in unlogged forests. The R^2^ is included to explain the fixed and random effects of the model. The F and P-values are the summary of the significance of the model.

### Merchantable volumes for all the timber species, mahogany species and other harvested tree species along the logging gradient

The volume per plot for all timber, mahogany, and other harvested species is provided in supplementary materials (S10). Overall, commercial timber volume was higher in unlogged forests than in logged forests (Table 7). GLMM results confirmed significantly greater timber volume in unlogged forests (β = 0.553, P = 0.022), while differences between lightly and heavily logged forests were not significant (β = 0.037, P = 0.786). Timber volume also varied significantly across plots (β = 0.156, P = 0.001).

**Table 7.** Merchantable volumes (m^3^/ha ± SD) for all timber trees, mahogany species and other harvested timber species in each of the logging intensity.

| Timber category | Diameter category (cm) | Heavily logged<sup>a</sup> | Lightly logged<sup>b</sup> | Unlogged<sup>c</sup> | F - Value | P - Value |
| --- | --- | --- | --- | --- | --- | --- |
| | | Volume ($\text{m}^3/\text{ha}$) | Volume ($\text{m}^3/\text{ha}$) | Volume ($\text{m}^3/\text{ha}$) | | |
| All timber species | $\geq 30$ | $45.43^a \pm 9.87$ | $66.81^{ba} \pm 23.39$ | $81.16^{ca} \pm 17.01$ | 14.49 | <0.001 |
| Mahogany species | | $5.08a \pm 4.78$ | $7.20b \pm 6.49$ | $25.79^{cb, ca} \pm 8.35$ | 26.78 | <0.001 |
| Other harvested species |  | 12.49<sup>a</sup> ± 8.59 | 20.99<sup>ba</sup> ± 12.48 | 31.89<sup>ca</sup> ± 14.21 | 8.15 | 0.001 |
| All timber species | ≥ 60 | 50.46<sup>a</sup> ± 16.86 | 82.38<sup>ba</sup> ± 55.28 | 130.22<sup>cb, ca</sup> ± 56.81 | 10.07 | <0.001 |
| Mahogany species |  | 3.53<sup>a</sup> ± 3.49 | 26.10<sup>ba</sup> ± 37.78 | 44.71<sup>ca</sup> ± 19.47 | 9.19 | <0.001 |
| Other harvested species |  | 27.78<sup>a</sup> ± 17.82 | 36.11<sup>b</sup> ± 31.21 | 55.89<sup>ca</sup> ± 22.52 | 3.09 | 0.056 |

Note: m<sup>3</sup> = Volume, ha = Hectare, SD = Standard deviation

GLMM results showed that mahogany species had significantly higher volume in unlogged forests compared to logged forests (β = 1.592, P = 0.001), with no significant difference between lightly and heavily logged forests (β = 0.407, P = 0.253). Similarly, other harvested timber species had significantly greater volume in unlogged forests (β = 0.846, P = 0.042), while differences between lightly and heavily logged forests were not significant (β = -0.259, P = 0.384). Timber volume also varied significantly across plots (β = 0.322, P = 0.008).

### Effect of time on timber volume recovery

We assessed the effect of time on timber volume recovery using plot-level data (S11, supplementary materials). GLM results showed a significant effect only for mahogany species, with no significant differences for all commercial timber species combined or other harvested species (Fig 5).

**Fig 5.**
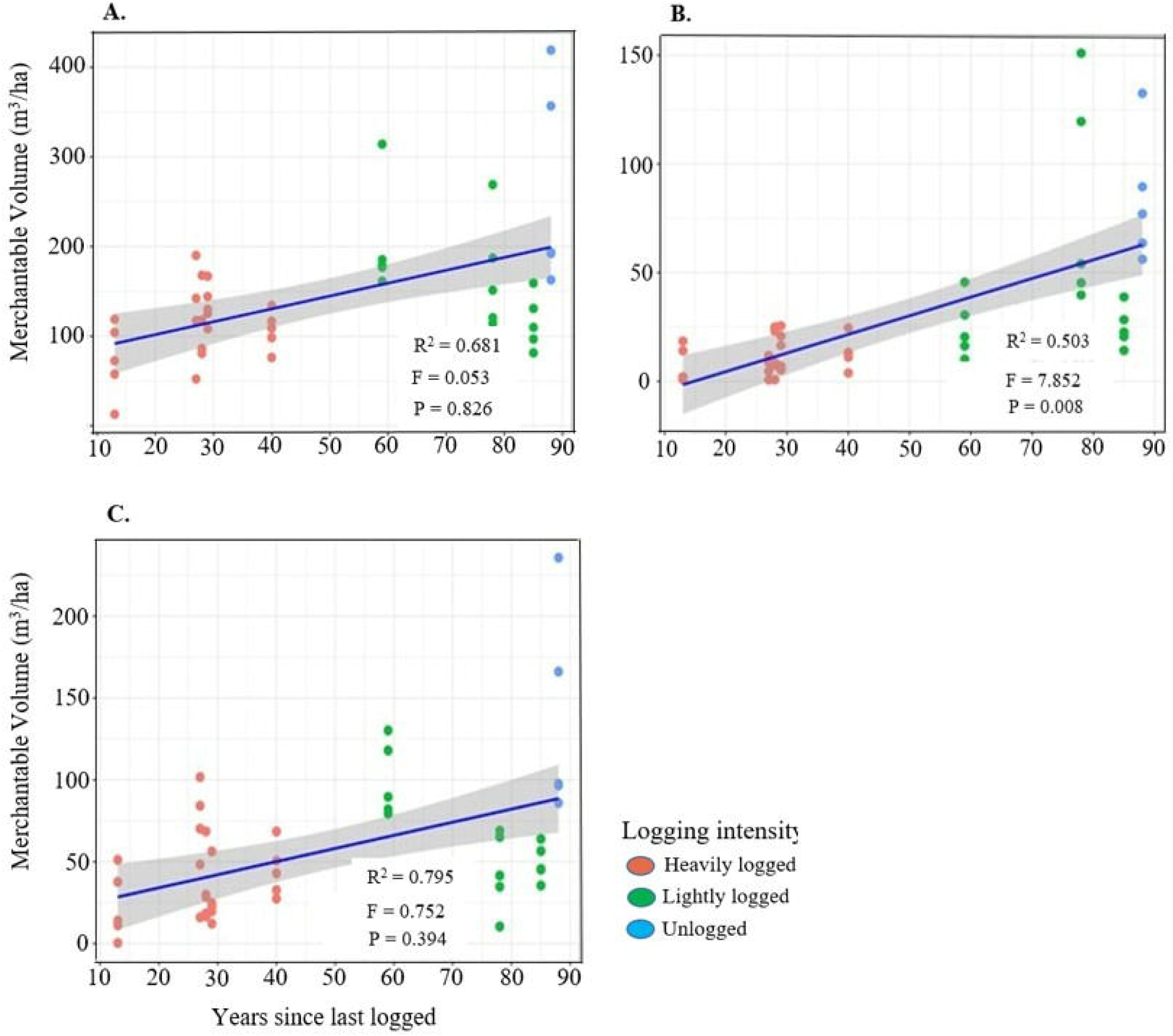
Merchantable/timber volume of (A) all commercial harvested timber trees, (B) mahogany species and (C) other harvested timber species against years the plots were last logged in Budongo Forest. Timber volume in each plot is represented by a single dot. Plots with **red** color represent heavily logged areas, **green** indicates lightly logged compartments, and **blue** corresponds to plots in unlogged forests. The R^2^ is included to explain the fixed and random effects of the model. The F and P-values are the summary of the significance of the model.

### Effect of enrichment planting and arboricide treatment on volume recovery of Mahogany (*Khaya anthotheca*) in Budongo forest

The effects of enrichment planting and natural regeneration on the volume of *Khaya anthotheca* are shown (Fig 5). Stand density analysis for trees with DBH ≥30 cm revealed that the enriched compartment (N1) had the highest density (28.8 ± 13.9 stems/ha), followed by the unlogged compartment (N15) (17.2 ± 4.5 stems/ha), while logged but not enriched compartments (B2: 11.6 ± 3.4 stems/ha; W20: 10.4 ± 5.1 stems/ha) had the lowest densities (S1**2**, supplementary material).

ANOVA indicated a significant difference among compartments (F₃, ₁₉ = 5.81, p = 0.008), with Tukey’s post hoc tests showing N1 had significantly higher densities than N15 (p = 0.034), B2 (p = 0.003), and W20 (p = 0.002), while differences between N15 and the logged compartments were not significant.

The merchantable volume of *K. anthotheca* extracted during logging and the volume recovered since the last logging event for each compartment are shown in Fig. 6A. The GLMM results indicate that timber volume recovery is significantly higher in enriched compartments compared to logged and naturally regenerated ones (Fig 6B). However, no significant difference was found between enriched and unlogged compartments, although some enriched plots had higher timber volumes. Arboricide treatment showed no significant effect on mahogany regeneration (β = -0.005, p = 0.986). Overall, the results suggest that enrichment planting boosts timber volume recovery, while arboricide treatment has little impact on mahogany regeneration.

**Fig 6.**
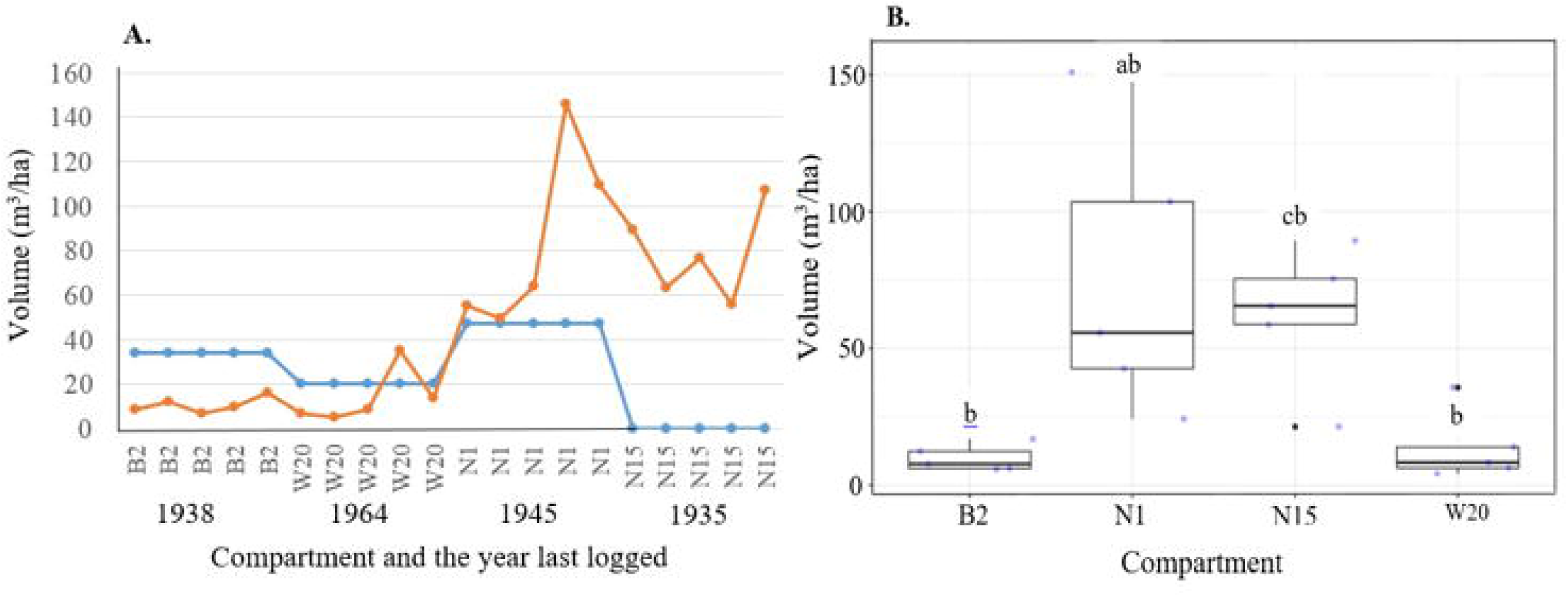
Comparison of volume harvested against volume accumulated over time since last harvesting (A) and merchantable volume (B) between enriched compartment with other compartments (logged and unlogged with only natural regeneration). Significance difference is shown using ^ab^ ^and^ ^cb^ symbol only.

## Discussion

### Forest recovery

Although our study lacks full replication, balance, and control for preexisting conditions, silvicultural treatments, and harvest intensities, the observed reduction in tree densities, with significant impacts on understory trees in both heavily and lightly logged forests, and on canopy trees only in heavily logged forests (Table 2 Appendix), reflects the historical trajectory of the forest. Specifically, we observed significantly lower stem densities of late-successional trees in logged forests compared to unlogged forests. Further analysis of the stem densities of four dominant late-successional species (*Drypetes ugandensis, Lasiodiscus pervillei, Celtis mildbraedii*, and *Rinorea beniensis*) in the undergrowth revealed that their density was lowest in heavily logged forests, followed by lightly logged forests, and highest in unlogged forests. This suggests that logging disrupts their recruitment and persistence. The general decline in stem densities of late successional species indicates that logging-induced-microhabitat changes may hinder forest recovery by favoring pioneer and light-demanding species, potentially altering successional trajectories and species composition. A 1940s study on Budongo Forest ecology documented distinct successional stages, showing the influence of past disturbances on current forest composition [67]. Recent studies highlight that logging disrupts succession, alters species composition in both the understory and canopy, and hampers forest recovery [64]. This impact is especially severe due to the removal of large timber species like mahogany, which damages remaining trees and causes high early-stage mortality [87]. Additionally, logging reduces seed production in key timber species such as *Khaya anthotheca* [88]; decreases seed tree availability [57], and disrupts regeneration patterns, indicating unsustainable harvesting practices [89].

Timber management in Budongo Forest began with the removal of large trees, focusing primarily on species like mahogany (*Khaya* and *Entandrophragma*) and other key species (*Cynometra, Milicia, Alstonia, Lovoa, Maesopsis, Erythrophleum, Mildbraediodendron, Morus,* and *Ricinodendron*) with DBH ≥100 cm. Older harvesting interventions were intensive in some compartments but targeted fewer trees, while recent salvage logging aimed at promoting uniform regeneration by targeting smaller trees (DBH ≥60 cm), resulting in more extensive tree removal per hectare. This practice, often conducted through pit sawing, was exacerbated by illegal harvesting, which reduced seed availability, disrupted natural regeneration, and caused additional damage to residual trees. The significantly lower stem densities for understory species and reduced basal area of canopy species in logged forests (Table 2) can be attributed to frequent illegal extraction of poles and timber in Budongo Forest. A study on illegally abandoned logs and cut poles revealed that 80% of the illegally harvested timber came from mahogany species (*Khaya anthotheca, Entandrophragma utile*, and *Entandrophragma angolense*), with other species, including *Albizia glaberrima, Albizia zygia, Maesopsis eminii,* and *Cordia millenii*, also illegally harvested [90]. Forest recovery may take actually long time depending on many factors as was observed in Uganda’s Kibale Forest, where selectively logged areas showed lower stem density and basal area even after 45 years The long-term effects are similar to those observed in Uganda’s Kibale Forest, where selectively logged areas showed lower stem density and basal area even after 45 years [46]. These illegal activities not only deplete timber resources but also disrupt the natural regeneration of both timber and understory species, delaying forest recovery. These findings highlight the lasting impacts of logging, particularly heavy and illegal logging, on forest structure and regeneration, further delaying recovery before the next harvesting cycle.

Tree recovery, assessed through stem density in understory and canopy species, is influenced by time since logging and varies depending on reference conditions and research objectives. Our findings show that time since logging significantly affects the recovery of small (DBH <10 cm), large (DBH ≥60 cm), and canopy trees, but has a limited effect on medium-sized (DBH 10–<60 cm) and understory trees (Fig. 2). This pattern likely results from differences in growth rates, recruitment success, competition, and species-specific responses to disturbances. Ideally, logging creates canopy gaps that enhance light availability, promoting the regeneration of small trees and light-demanding pioneers, facilitating their transition into medium-sized trees [26](. However, medium-sized and understory trees showed limited recovery, suggesting that past logging intensity, environmental conditions, and ongoing human activities, such as selective harvesting and pole cutting, continue to influence recovery. This limited regeneration may indicate recruitment bottlenecks, competition, or growth suppression, slowing progression into larger size classes and delaying overall forest recovery. These findings align with other studies for example, in Sabah, Malaysian Borneo, [74] found that logging negatively impacted trees in the 10–40 cm and ≥60 cm DBH size classes even 23–35 years post-disturbance. Similarly, in Sarawak, Malaysian Borneo, [91] observed that dipterocarp tree recovery was more pronounced in tree with diameter (DBH ≥50 cm), while stem recovery remained low for smaller size classes (<50 cm) even 10–20 years after logging, indicating limited regeneration. In Budongo Forest, medium-sized tree recovery is hindered by ongoing pole cutting for sawing platforms, house construction, and tool handle production, with few stems in the target size class (DBH ≥5–14.9 cm). Additionally, limited colonization by pioneer species, such *as Maesopsis eminii, Macaranga* spp*., and Albizia* spp., further constrains recovery, with no significant variation in abundance across logging intensities. This is likely due to canopy closure over an extended post-logging period (13–85 years), which reduces light availability and suppresses pioneer recruitment. In contrast, studies conducted shortly after logging (1–5 years) showed open gaps favoring pioneer species establishment [91,92]. As canopy gaps close over time, vertical light gradients intensify, favouring shade-tolerant, late-successional species over pioneers [93], a trend also evident in our findings from Budongo Forest.

Our findings indicate that both mean tree heights and canopy heights (Table 5) were lower in logged forests compared to unlogged forests, emphasizing the long-term impacts of large tree removal on forest structure. In Budongo, the majority of canopy trees are key timber species, and unsustainable logging practices, combined with illegal harvesting of many canopy species [90], likely contribute to reduced canopy tree densities, thereby potentially hindering structural recovery. Other studies have shown that structural recovery in logged forests is a slow process influenced by logging intensity and the capacity of remaining tree species to utilize available resources for regeneration [25]. For instance, a study in Pasoh Forest, Malaysia, found that logging significantly reduced canopy heights even 53 years post-harvest [94]. Similarly, research in Sabah, Malaysian Borneo, using airborne LiDAR, revealed that heavily and moderately logged forests had reduced forest structure compared to old-growth plots, indicating that canopy recovery can take decades [28]. Excessive open space also affects vertical tree growth, as demonstrated by studies showing that increased light availability and reduced large-tree density cause trees near forest edges and in fragmented forests to be 30–40% shorter than those in closed forests [95]. This effect may be further exacerbated by frequent illegal logging, which disrupts structural recovery [96].

### Regeneration patterns of selected/harvested commercial timber species and abundance of tree species

Our findings showed poor regeneration of most commercial timber species with exception of *K. anthotheca* in class I that had fair regeneration and *C. alexandri* in class II with good regeneration in all the logging intensities (Table 6). The majority of timber species had juvenile populations of less than 1 stem/ha, indicating poor regeneration patterns, likely due to the low population of mature seed-bearing trees essential for seed production. For instance, *Olea capensis* (Class I) and *Ricinodendron heudelotii* (Class II) exhibited no regeneration in heavily logged forests. This phenomenon may be attributed to insufficient seed production, seed predation, or dispersal limitations. In Budongo Forest, tree regeneration failure has been primarily attributed to the effects of logging on microclimatic conditions, the reduction of seed-bearing trees[57], and recent findings on *Khaya anthotheca*, which demonstrated that higher adult conspecific abundance in the canopy enhances regeneration [97]. Elsewhere, a study in Borneo found that higher tree basal area was linked to increased seedling density, highlighting the importance of preserving large canopy trees for sustaining fruit production and promoting germination rates in tropical forests [98]. Thus, it is reasonable to attribute low regeneration of timber species in Budongo Forest to overharvesting of large trees as has been observed in Central Amazon [99]. Other studies in other tropical forests equally found low regeneration of timber species, for instance, a study in Tres Garantías, Mexico, found that *Swietenia macrophylla* (mahogany) juveniles were too scarce and insufficient to sustain future harvesting rates [100]. Again caution should be taken while interpreting high adult populations to increase in juvenile regenerations because some species like *Maesopsis eminii*-long-lived pioneer species had higher adult populations in lightly and unlogged forests but relatively low juvenile populations compared to the trend exhibited in heavily logged forests. We also noticed poor regeneration of *Entandrophragma* species (*E. cylindricum, E. Utile* and *E. angolense*) in Budongo forest in line with a study by [101] that reported slow recovery of Entandrophragma species across tropical forests in Africa. As noted by [102], juveniles are more sensitive to environmental factors than adults of the same species and tree species may respond differently to the microclimatic conditions created by logging disturbance. Poor recovery of juveniles of many tree species in logged forests has been linked to functional traits, as high-intensity logging tends to favor species with higher specific leaf area and low wood density that are mostly pioneer species [103,104]. Importantly, other factors beyond logging intensity influence tree and timber species recovery in Budongo and other logged tropical forests. Regeneration may be suppressed by inadequate propagules, seed dispersal limitations, seedling’s vigour, predation, intraspecific or interspecific competition [105,106]; High-frequency and conventional logging increase disturbance, degrade residual stand quality, and hinder seedling growth, reducing forest resilience [33,107]; Natural regeneration relies on recovery time, residual growth, and seed dispersal, while silvicultural interventions like enrichment planting boost recruitment [107]; Low soil nutrient levels and insufficient moisture, with nutrient limitation varying by soil type and degradation, hinder seedling growth [108]; Unregulated logging [49,96] and climate stressors [69] further hinder regeneration by disrupting natural processes, reducing habitat quality, and altering species composition, all of which impede recovery and species establishment.

Our findings revealed a low proportion of commercial Class I timber species in the medium size category (≥30 to <60 cm) in both heavily and lightly logged forests (Fig. 3), suggesting future timber volume from these species may be insufficient for upcoming harvest cycles. Larger Class I trees (DBH ≥60 cm), recommended for harvesting, were scarce. No *Entandrophragma* spp., *Milicia excelsa*, or *Lovoa trichilioides* individuals ≥60 cm were found in heavily logged areas; lightly logged forests had only one *Entandrophragma cylindricum*, three *Milicia excelsa*, and no *Lovoa trichilioides.* The density of *Khaya anthotheca* stems ≥60 cm was limited, with 2.53 stems/ha in heavily logged, 8.53 stems/ha in lightly logged (mainly in enriched compartment), and 7.69 stems/ha in unlogged forests. This low abundance of Class I species is likely due to intensive logging, which hinders regeneration and poses a risk of reproductive isolation and population decline in logged areas. Overharvesting has driven the decline of American mahogany (*Swietenia macrophylla*) in Brazil, Peru, Bolivia, and Colombia [109]. While low-intensity logging can aid timber stock recovery, it requires strict regulations on cutting limits, residual stocking, and harvest restrictions [110]. However, single-tree selection may not create sufficient canopy gaps for light-demanding species like mahogany [111]. Our findings suggest that Class II commercial species (Fig 2) could serve as an alternative timber source in future harvests if well managed. A similar trend was observed in the Brazilian Amazon, where 32 years after the first cut, timber volume had not fully recovered, necessitating the inclusion of previously unlogged species for future harvests [112]. To ensure long-term sustainability, promoting the regeneration of commercially valuable species through selective management and enrichment planting is crucial.

### Effect of forest structural parameters (basal area, stem density and canopy cover) on regeneration status/juvenile abundance of all timber species, mahogany species and other harvested timber species as mediated by logging intensity

From our results, only basal area and unlogged forest had significant effects on abundance (stems/ha) of mahogany juveniles (Table 4). Our results showed significant positive interactions between basal area and lightly logged or unlogged forests on mahogany juvenile abundance, suggesting that the presence of large seed trees in these areas may support mahogany regeneration. Overall, juvenile abundance decreased with basal area, likely due to competition avoidance, particularly for light. Again as noticed from our seed trap experiment (personal observation), seeds of mahogany was able to reach some traps even in plots were the big/mother trees were never available. Implying, big mahogany trees tend to disperse a lot of seeds far away as a strategy to reduce the effect of density-dependent on the survival of new recruits. A recent study in Budongo Forest found that juveniles of mahogany (*Khaya anthotheca*) were more abundant farther from the mother trees, suggesting a strategy to avoid conspecific competition [113].

Higher seedlings density was also found in unlogged forest compared to logged forest due to possibly high density of adult trees of the focus species [114]. Conversely, the negative correlation between stem density and mahogany juvenile abundance in unlogged forests indicates possible high interspecific competition for resources or negative distance-dependent effects from intraspecific competition and attacks by host-specific natural enemies [115]. Greater competition from pre-existing plants in the unlogged areas and also to greater extent competition from weeds in the felling gaps affects juvenile abundance [116]. Most studies on distance-dependent effects have focused on pathogens and insect herbivores [117], but mammals may also play a key role in shaping plant recruitment in tropical forests [118]. Our preliminary analysis of seed removal in Budongo Forest revealed that mahogany seeds are highly removed by seed predators across all compartments, suggesting a strong negative impact on regeneration in areas with low seed production.

The significant effect of time on commercial timber juvenile abundance recovery (stems/ha) (Fig 5) suggests that as years progress, forest conditions increasingly favor the recruitment of late-succession or shade-tolerant species, such as mahogany. The removal of large, crown-spreading trees, particularly mahogany, alters understory and canopy structures [119], This change initially supports the establishment of light-demanding species-mostly pioneers over shade-tolerant species-late succession species. However, for mahogany species (*Swietenia macrophylla*) and a few other late-succession species, greater disturbance (such as intensive canopy opening) has been shown to support their regeneration [120]. Our results (Table 5) showed the lowest average canopy cover (75.9%) in heavily logged forests, with canopy cover not significantly affecting mahogany regeneration. However, the strong negative correlation of canopy cover with juvenile abundance in unlogged areas (β = -0.976) compared to lightly logged areas (β = -0.068) suggests that areas with lower canopy cover (more openings) favor juvenile abundance of commercial timber species. This trend is also evident in the variability of juvenile abundance across plots (Fig 5) with different logging intensities for commercial timber species, including mahogany and other harvested species, particularly in heavily logged forests. This raise the issue that tropical forests are constantly changing [121] and thus, disturbance such as logging could be increasing the pace of change process.

### Effect of logging intensity on merchantable volume recovery of commercial timber species

Our results showed that logging significantly impacted timber volume recovery for all categories i.e. all commercial timber species combined, including mahogany and other harvested species, with unlogged forests having the highest timber volumes and thus, supporting our second hypothesis. Also plot timber volumes varied significantly among and across logging intensities. The reason is that commercial logging in Budongo forest-Uganda extracted higher timber volumes i.e. 39.9 (range 25.6 – 58.7) m^3^/ha from 13 commercial species in most of the compartments. The timber volume extraction for mahogany species only i.e. 28.5 (16.3 – 47.1) m^3^/ha constituted approximately 71.4% of the overall timber volume extracted from all the 13 timber species. This average volume off-take of approximately 40 m^3^/ha for all the commercial species and 29% for mahogany species in Budongo is far above most of the recommended volume off-takes for sustainable logging in tropical forests [49]. Such high volume extraction coupled with ineffective post-harvest silvicultural treatments affect timber volume recovery since most tropical timber species have been shown to have very slow growth rates [34,122,123]; with exception of a few long-lived pioneer species with high juvenile selection effect that allows them to grow very fast to recover significant timber volume over time [122,124].

Even after 60 years had elapsed, lightly logged forests/compartments had not recovered in terms of timber volumes for mahogany and other harvested timber species to the level closed to the unlogged forest, supporting our second hypothesis. The significant variation in plot timber volumes could be due to differences in species distribution and or the low timber volumes could be as well a result of frequent illegal logging of highly targeted timber species (Personal observation). In Budongo forest, illegal logging if not properly handled will continue to affect the timber volume recovery of desired timber species. Again, high intensity logging has been shown to affect volume recovery of timber species even over long period of time[22,107]. For example, in the tropical forests of French Guiana and Paracou, a logging intensity of 8 – 29 m^3^/ha was modelled for timber recovery, showing that approximately 4.1 m^3^/ha and 6 m^3^/ha of timber were recovered after 65 years, respectively [22]. In Amazon, a systematic review study revealed that high intensity logging of 20 – 40 m^3^/ha affected timber volume recovery for the next logging cycles [32]. Precisely, where logging policies and forest protection laws are not well implemented like the case in Uganda [2], recovery of timber volumes for the next logging cycles is most likely not attainable.

We tested the effect of time on timber volume recovery, and our results showed a significant difference only for mahogany species, not for all timber species combined or other harvested species (Fig 5). This may be due to an insufficient number of residual and reserved “seed trees” left during mechanical logging in the 1990s, along with illegal logging of mahogany in many compartments. Additionally, re-logging of some compartments from 1990 to 2010 through pitsawing likely reduced the stem densities of retained mahogany “seed trees” and large residual trees, further impacting volume recovery. Residual trees play a crucial role in timber volume recovery in logged forests [23]. In Budongo, the recovery of species and timber volumes in logged forests was impacted even after 60 years due to an insufficient number of retained “seed trees” to support sustainable passive restoration [125]. While natural regeneration from seeds or sprouts is faster and more cost-effective than active regeneration (planting of seedlings), successful seed regeneration depends on sufficient seed production, effective dispersal, and a suitable environment for juvenile germination and establishment [126].

### Effect of enrichment planting and arboricide treatment on volume recovery of mahogany species (*Khaya anthotheca*) between compartments logged 60 years

Our result found a striking difference over a period of 60 years whereby compartment which was the most heavily logged in terms of timber volume per heactare but was properly enriched in 1950s had recovered significantly higher timber volume of *K. anthotheca* compared to the other two logged but not enriched compartments (Table 7 and Fig 6). Arboricide treatment had no effect on the volume of mahogany recovered in the compartments where it was applied. Therefore, it is evident that the increase in timber volume after silvicultural treatment was due to successful establishment and higher stem density of *K. anthotheca* tress with (DBH ≥30 cm) in enriched compartment (N1) compared to the other natuarally regenerated logged compartments (B2 and W20). We found natural regeneration alone had not enhanced the volume recovery of *K. anthotheca* logged forests to the level of unlogged forest after over 60 years had elapsed and thus, rejecting the third hypothesis. However, within the same period with increased protection offered to enriched (N1) and unlogged (N15) compartments, the timber volume of *K. anthotheca* in enriched compartment had recovered to the level of unlogged forest compartment (N15), supporting our third sub-hypothesis. Our findings on successful mahogany enrichment planting in logged compartment of Budongo Forest align with global studies. In Cameroon, *Pericopsis elata* enrichment planting achieved 61% survival after five years, even without maintenance [127]. Similarly, in Pará, Brazil, *Swietenia macrophylla* (mahogany) had 61% seedling survival after 4.4 years, indicating potential gains in timber volume and profitability [128]. Enrichment planting combined with tending treatments was the most effective strategy for recovering commercial species where natural regeneration was limited [129]. Additionally, planting in larger gaps outperformed smaller gaps, likely due to improved light conditions for early-stage timber growth [130]. In Borneo, line planting of dipterocarp species in twice-logged forests restored timber stocks to primary forest levels within 40 years, whereas natural regeneration achieved less than half that volume in 60 years [131]. Similarly, sapling and pole stem density increased over time in actively restored areas, while naturally regenerating forests had higher initial density but showed no further increase with time [20]. Therefore, it reasonable to conclude that, heavily logged forests if left to regenerate naturally without effective protection, enough retention of seed trees and other silvicultural practices like enrichment planting, climber cuttings among others may fail to significantly recover timber volumes of highly targeted species for the next harvesting cycles.

Much as enrichment planting in Budongo Forest was abandoned in 1960s due to high establishment costs and mortality rates caused by mostly elephants, Buffalos, wild pigs and bushbucks [56], our result of increased timber volume production or recovery by enrichment planting for next logging cycle adds to already attested contributions of active restoration such as promotion of successful regrowth, enhancement of forest structure, increasing species composition and economic value of logged forests over time [54,132,133]. Although we did not assess timber profitability, our findings suggest that if financial conditions are favorable, enrichment planting could bolster the economic value of production forests [134]. While our results, along with other studies [135,136], highlight its benefits, it is also important to acknowledge the challenges that need to be addressed. These include the simplification of species richness and diversity, reduced genetic diversity due to limited mother seed sources, high costs, and poor establishment from high seedling mortality [98,137]. Addressing these issues is essential for active restoration to sustainably enhance timber production in logged forests [138]. A limitation of our study is that we did not compare *K. anthotheca* tree quality (specifically tree architecture) between enriched and non-enriched compartments which would have provided further insights into the impact of enrichment planting on timber quality.

### Implications for biodiversity conservation and sustainable timber production

Timber production in species rich forests, such as Budongo, can be managed sustainably to prevent regeneration failure [139]. Globally, 25% of remaining primary tropical forests are designated for timber extraction. With timber demand expected to rise by 30% by 2050 [5] and widespread illegal logging [96], sustainable management is essential, as biodiversity conservation cannot rely solely on protected areas. Thus, sustainable logging at lower intensities, using best practices like Reduced Impact Logging [140], can help maintain ecological functions, while well-managed tropical production forests have been shown to support high biodiversity [14,29]. Implying, the sustainability of logged forests for timber and biodiversity depends on understanding logging impacts, species recovery, and management effectiveness, including enrichment planting and climber cutting. Therefore, as governments, including Uganda, strengthen commitments to the Aichi Biodiversity Targets and the Paris Agreement on climate change, uncertainty remains about the role of improved forest stewardship in achieving climate mitigation and biodiversity conservation goals [141]. Research on forest structure, timber regeneration, and volume recovery in logged forests provides crucial insights into biodiversity responses and informs sustainable forestry practices [18,21,52]. However, in Uganda, such research remains limited due to challenges in long-term biodiversity monitoring and the absence of pre-logging data, as most tropical forests were logged between the 1940s and 1960s and the need to conduct research to provide information for enhanced sustainable management of logged tropical forests. Our findings indicate that logging affects the regeneration of key timber species and timber volume recovery even after 60 years had elapsed, highlighting the need for targeted management to enhance timber species regeneration. Again in Budongo Forest, passive natural regeneration alone has not supported the recovery of high-value timber species like mahogany (*Khaya anthotheca*), whereas enrichment planting has significantly enhanced mahogany timber volume in heavily logged compartments. Similarly, studies in the Amazon and Southeast Asia show that actively restored forests can recover timber stocks comparable to primary forests within decades, while passive regeneration often results in lower commercial species abundance [45,74,128,131]. These findings emphasise the need for science-based silvicultural interventions to ensure that logged forests in Budongo sustain both ecological and economic benefits. We recommend enrichment planting with economically desirable timber species such as *Khaya anthotheca*, *Entandrophragma* spp., Lovoa spp., and *Milicia excelsa*, alongside fast-growing species like *Maesopsis eminii*., *Albizia* spp. Additionally, integrating this approach with sustainable harvesting practices, such as Reduced Impact Logging and broader species selection from classes II, and III, can enhance timber yields while conserving biodiversity in production forests.

## Conclusions

Our results implied that logging and associated differences across the forest affected the implied recovery of forest structure, commercial timber species regeneration and volumes. The high abundance of small stem diameters in unlogged compared to logged forests could be attributed to adequate regeneration of tree species due to high populations of adults. Overall, canopy cover had partially recovered in even heavily logged forest after just a period of less than 30 years and high intensity logging has effect on timber volume recovery especially of high value and frequently harvested timber species such as mahogany species. Noticeably, passive regeneration has been insufficient to permit full recovery of both density and timber volumes of highly targeted species compared to active regeneration (enrichment planting) as evident in our findings on recovery of *K. anthotheca* in compartments logged before 1964. If the costs can be justified we recommend enrichment planting to enhance timber species and volume recovery, reduced logging intensity to less than < 15 m^3^/ha as demonstrated by other studies in the Amazon [18] and Asia [17].

## Appendix

**Table 2.**
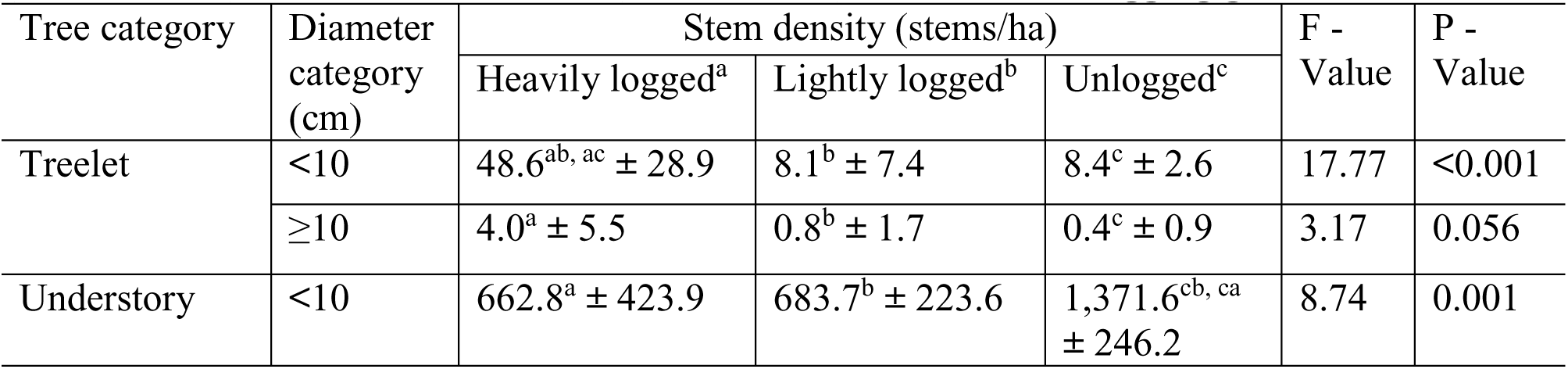
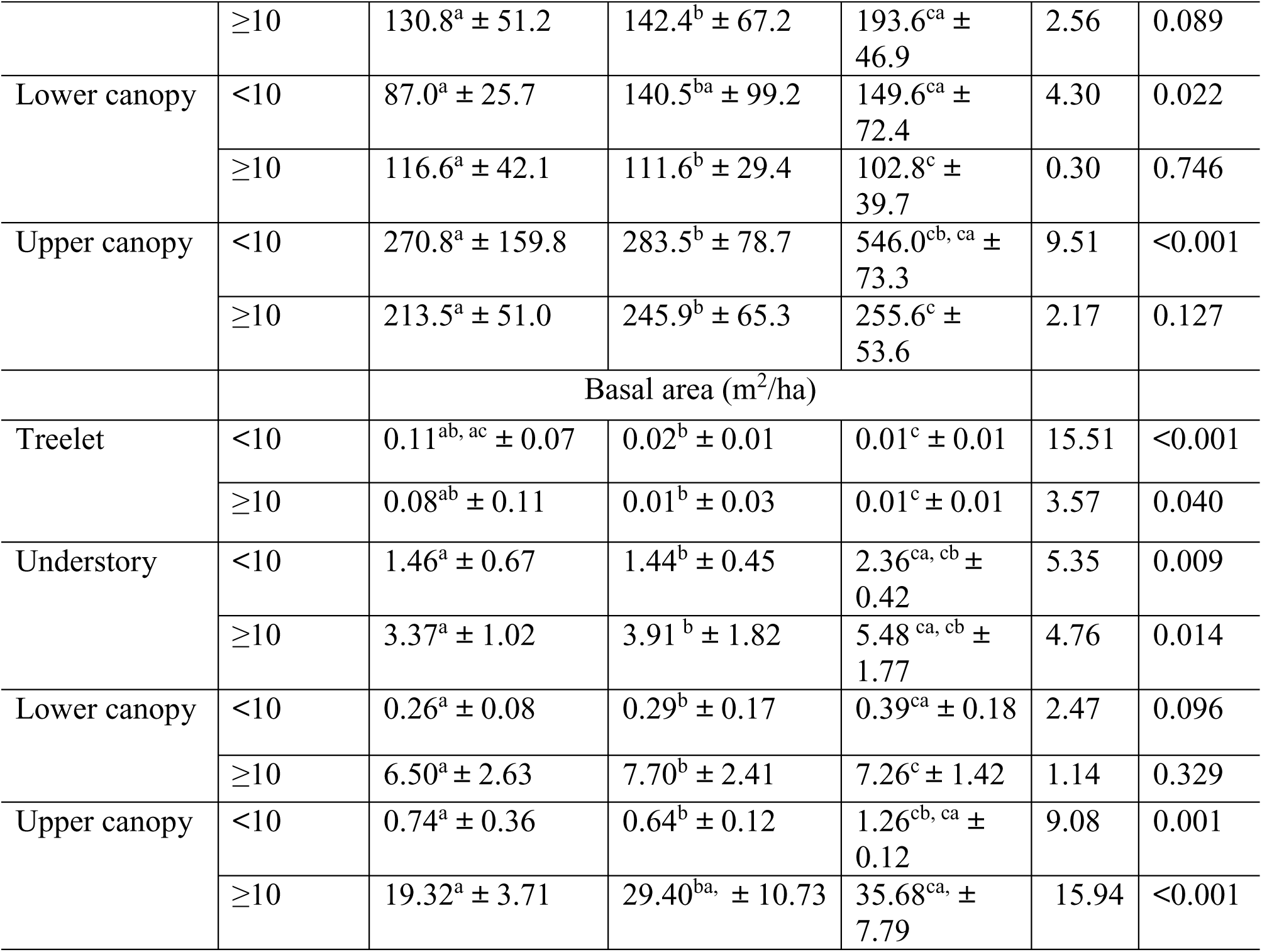
Summary of forest structure i.e. stem density (stems/ha ± SD) and basal area (m^2^/ha ± SD) for vertical stratification of tree growth-form for diameter categories of small trees (DBH <10 cm) and established trees (DBH ≥10 cm) in each of the logging gradient.

## Supporting information

S1 Table. Dataset used for the analyses of total tree counts for each categories along logging intensities in Budongo Forest.

S2 Table. Dataset used for calculating forest structure i.e. stem density and basal area for vertical stratification of tree growth-forms along logging intensities in Budongo Forest.

S3 Table. Dataset used for calculating the effect time since logging on stem density of small, medium, large, understory, and canopy trees in Budongo Forest.

S4 Table. Dataset used for calculating the stem density and basal area of all timber, mahogany, and other harvested species in Budongo Forest.

S5 Table. Dataset on succession types across logging intensities in Budongo Forest.

S6 Table. Dataset used for calculating variation in forest structure i.e. mean stem diameter, stand dominant heights, canopy closure, and undergrowth along logging intensities in Budongo Forest.

S7 Table. Dataset used for calculating regeneration status of key timber species harvested in Budongo Forest.

S8 Table. Dataset used for calculating the proportions of class I_II and III across logging intensities in Budongo Forest.

S9 Table. Dataset used for calculating the effect of forest structural parameters on Juvenile abundance of all timber species, mahogany and other harvested species in Budongo Forest.

S10 Table. Dataset used for calculating the merchantable volume for medium and large diameter trees of all timber, mahogany and other harvested species in Budongo Forest.

S11 Table. Dataset for calculating the effect of time on merchantable volume of all timber, mahogany, and other harvested species in Budongo Forest.

S12 Table. Dataset used for calculating the effect of enrichment planting and arboricide application on the recovery of Khaya anthotheca volumes in Budongo Forest.

## Acknowledgements

The field research was supported by Makerere University-Kampala, National Forestry Authority (NFA) and Budongo Conservation Field Station (BCFS). We deeply appreciate the support of Parabotanists (Obed Goffin, Nelson Adaku, Micheal Ezati, and Isaac Moko) from BCFS and other casual labourers (Kennedy Pithua, Jackson Okuti, Charles Drani, Emmanuel Toko and Feb Oloya) who helped with data collection. The authors would like to thank in a special way the independent reviewers for their valuable comments that improved the manuscript.

## Author Contributions

Conceptualisation: David Kissa Ocama, Emmanuel Fred Nzunda, Mnason Tweheyo, Daniel Lussetti, Douglas Sheil

Data curation: David Kissa Ocama

Formal analysis: David Kissa Ocama

Funding acquisition: David Kissa Ocama

Investigation: David Kissa Ocama

Methods: David Kissa Ocama, Emmanuel Fred Nzunda, Mnason Tweheyo, Daniel Lussetti, Enock Ssekuubwa, Douglas Sheil

Software: David Kissa Ocama, Enock Ssekuubwa

Supervision: Emmanuel Fred Nzunda, Mnason Tweheyo, Daniel Lussetti

Validation: Emmanuel Fred Nzunda

Visualization: David Kissa Ocama, Mnason Tweheyo, Daniel Lussetti, Douglas Sheil

Writing – original draft: David Kissa Ocama

Writing – review & editing: David Kissa Ocama, Emmanuel Fred Nzunda, Mnason Tweheyo, Daniel Lussetti, Enock Ssekuubwa, Douglas Sheil

